# MAP4-MAP7D1 partitioning on tyrosinated-detyrosinated microtubules coordinates lysosome positioning in nutrient signalling

**DOI:** 10.1101/2025.10.07.680844

**Authors:** Deepak M. Khushalani, Joydipta Kar, Satya Bikash Nayak, Subhash Chandra Chaudhary, Nitin Mohan

## Abstract

Microtubule-associated proteins (MAPs) and tubulin post-translational modifications (PTMs) together shape a dynamic intracellular landscape for motor-driven transport, yet how the “MAP-PTM crosstalk” regulates organelle positioning remains unclear. Here, we show that MAP4 and MAP7D1 selectively partition onto distinct microtubule subsets demarcated by tyrosination and detyrosination, respectively, creating specialized tracks for kinesin motors. MAP4’s preferential binding depends on its projection domain, while expanded microtubule lattice states mediate MAP7D1’s enrichment on detyrosinated microtubules. Remarkably, rigor kinesin-1 (KIF5B-R) predominantly localizes to detyrosinated, MAP7D1-coated tracks, whereas rigor kinesin-3 (KIF1A) prefers tyrosinated, MAP4-decorated microtubules. We further find that the local density of MAP4 and MAP7D1 on microtubules fine-tunes lysosomal movement and directional transport. Moreover, MAP density is modulated to coordinate lysosomal reorganization in response to nutrient availability. During starvation MAP7D1 density on microtubules increases while MAP4 density decreases, localizing lysosomes to the perinuclear region. Conversely, with nutrient stimulation, MAPD1 density declines, allowing lysosomes to migrate towards the cell periphery. Altering the cellular levels of MAP4 and MAP7D1, either up or down, hinders lysosomal motility, trapping them near the nucleus and impairing their responsiveness to nutrient stimulation. Together, our findings reveal two distinct MAP–PTM circuits, a MAP4–tyrosination–kinesin-3 axis and a MAP7D1– detyrosination–kinesin-1 axis, that govern lysosome positioning for nutrient signaling, highlighting the combinatorial logic of MAP and tubulin codes in shaping microtubule function.

## Introduction

Microtubules, polymerized from α/β-tubulin heterodimers, form a polarized and dynamic cytoskeletal network extending from the microtubule-organizing center (MTOC) to the cell periphery. Beyond facilitating cell shape, cell division, and mechanotransduction, microtubules serve as tracks for the directed transport of intracellular organelles. Lysosomes leverage this network to rapidly reposition in response to metabolic cues^1^. During nutrient deprivation, lysosomes accumulate near the nucleus to support autophagy, whereas, in nutrient-rich conditions, they redistribute to the cell periphery to enable mTORC1 activation —a master regulator of cell growth and metabolism^2–4^. This dynamic repositioning of lysosomes is mediated by motor proteins: dynein drives retrograde movement towards the MTOC, while kinesins enable anterograde transport to the cell periphery^5–8^.

Recruitment of kinesin motors to lysosomes is governed by lipid and protein cues on the lysosomal surface. Nutrient availability triggers a phosphoinositide switch from PI4P to PI3P, enabling the recruitment of KIF16B, a kinesin-3 family motor, to drive lysosomes to the periphery^9^. Small GTPases, such as Arl8b, RILP, and the BORC complex, also link kinesin-3 motors like KIF1A and KIF1Bβ to lysosomes, enabling peripheral lysosomal movement crucial for efficient mTORC1 activation^10–12^. While much is known about the lipid and protein machinery on the lysosome surface that primes them for transport, how they integrate with spatial cues on the microtubule surface remains unclear. Microtubules are not uniform—they are marked by diverse microtubule-associated proteins (MAPs) and post-translational modifications (PTMs) that generate a variable and dynamic surface landscape^13,14^. How these molecular signatures on the microtubule surface are spatially organized and coordinate with the lysosome surface machinery to enable the rapid reorganization of lysosomes in response to cellular cues remains unclear. Tubulin proteins undergo a range of PTMs such as acetylation^15^, detyrosination^16,17^, tyrosination^18,19^, polyglutamylation^20,21^, and polyglycylation^22^—that define distinct subpopulations of microtubule tracks. Motor proteins differentially interpret these PTMs—the Tubulin code^13^. Kinesin-1, for example, preferentially localize to detyrosinated or acetylated microtubules^6,23–25^, while kinesin-3 motors prefer tyrosinated tracks^6,26^. Interestingly, kinesin-1 motility is impaired on detyrosinated microtubules, resulting in an enrichment of lysosomes on detyrosinated microtubules, facilitating their fusion with autophagosomes^27^. In parallel, MAPs dynamically bind to microtubules and selectively tune motor behavior —the MAP code^14,28^. For instance, MAP7 promotes kinesin-1 recruitment and motility but inhibits kinesin-3 and dynein^29–31^, whereas Tau-family MAPs (Tau, MAP2, MAP4) generally suppress kinesin-1 driven transport^32–35^.

Emerging evidence suggests that PTMs and MAPs do not operate independently but engage in complex regulatory crosstalk. For instance, MAP4 restricts access to the detyrosination enzyme, vasohibin to the microtubule *in vitro*^36^, while MAP7 promotes αTAT1-driven acetylation by stabilizing expanded microtubule lattices^37^. Minus-end binding proteins such as CAMSAP2/3 also promote lattice expansion and increase detyrosination^38^. Moreover, MAPs show preferential associations with specific microtubule subsets—MAP7 with acetylated^37,39^, DCLK1 with dynamic microtubules^26^, and MAP4 with both tyrosinated and detyrosinated microtubules^40^. Building on these findings, we hypothesize a potential MAP-PTM code that may regulate motor activity to facilitate lysosomal repositioning and function.

Here, using super-resolution microscopy, we uncover distinct associations between epithelial MAPs and modified microtubule subsets. We find that MAP4 preferentially localizes to tyrosinated microtubules via its projection domain, while MAP7 is enriched on detyrosinated lattices through an expanded microtubule conformation. Kinesin-3 (KIF1A) and kinesin-1 (KIF5B) motors associate with MAP4- and MAP7D1-decorated tracks, respectively. High-density MAP4 and MAP7D1 impair anterograde lysosomal motility more than retrograde movement, restricting lysosome dispersion and attenuating lysosomal remodelling in response to nutrients. Our findings identify MAP4 decoration on tyrosinated microtubules and MAP7D1 enrichment on detyrosinated microtubules as key regulators of kinesin-3 and kinesin-1mediated lysosomal positioning and function.

## Results

### MAP4 and MAP7D1 exhibit largely non-overlapping localization on the microtubule network

To examine how distinct MAPs are spatially organized along the microtubule network, we focused on two prominent epithelial MAPs—MAP4 and MAP7D1, the predominant microtubule-bound MAP7 isoform in BS-C-1 cells (Figure S1a). Confocal imaging revealed microtubule strands that are distinctly decorated by either MAP4 or MAP7D1, and a sub-population co-decorated by both MAPs (Figure 1a). To rule out the possibility that the apparent non-overlap of MAP4 and MAP7D1 arises from antibody occlusion during immunostaining, we performed sequential staining of MAP4 followed by MAP7D1 and vice versa (Figure 1a and S1b). Both approaches yielded consistent and distinct spatial distributions, confirming that MAP4 and MAP7D1 are largely non-overlapping.

**Figure 1:**
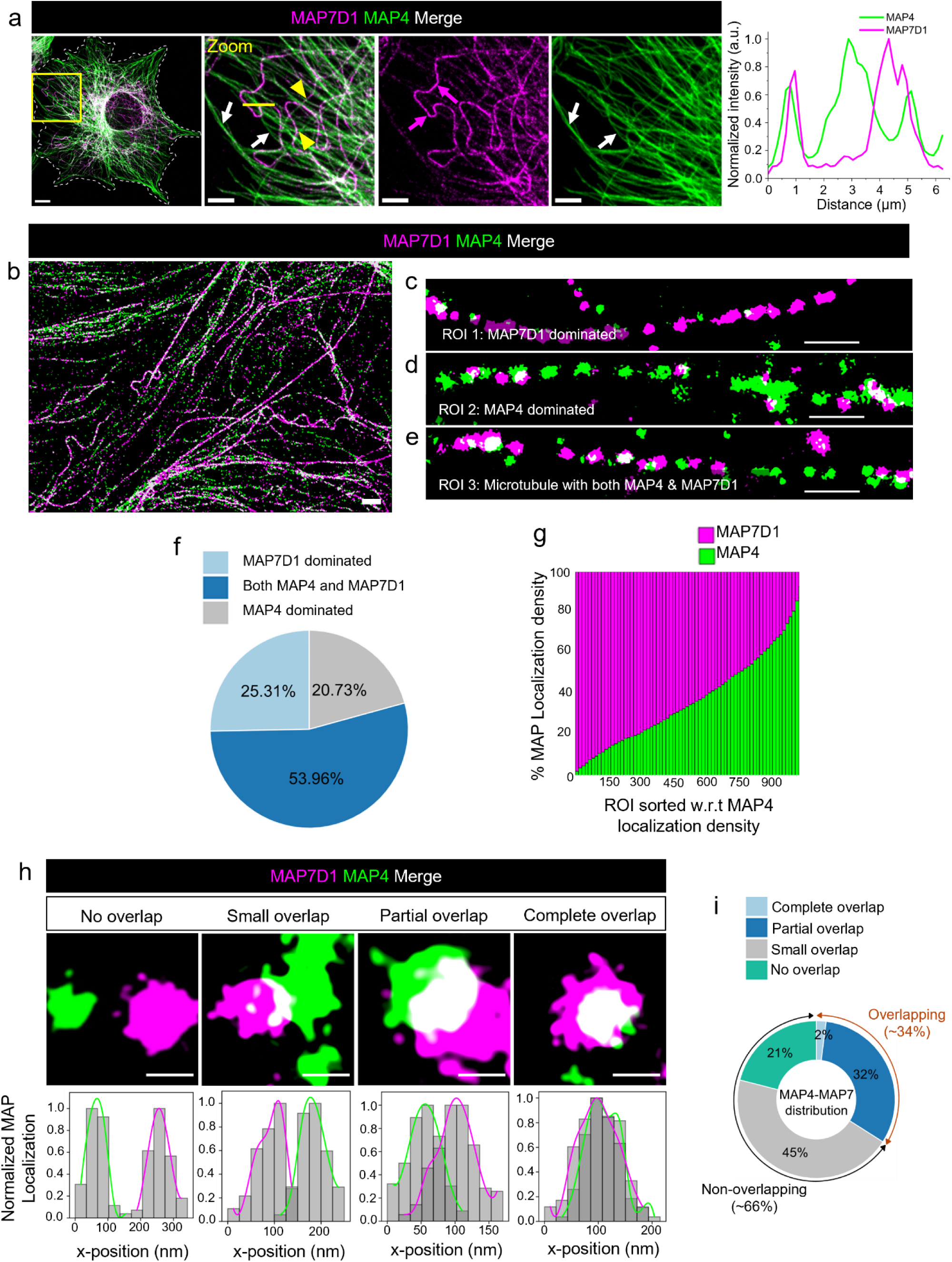
MAP7D1 and MAP4 nanoclusters are largely exclusive on microtubule network. (a) Dual-colour confocal images of BS-C-1 cells showing the distribution of microtubule-bound MAP7D1 (magenta) and MAP4 (green). Magenta arrows indicating MAP7D1-dominated microtubules, white arrows indicating MAP4-dominated microtubules, and yellow arrowheads marking microtubules decorated by both MAPs. A corresponding line intensity plot along the yellow line in the zoomed image reveals that the two MAPs are predominantly associated with distinct microtubule tracks. (b) Two-colour STORM image showing the nanoscale distribution of MAP7D1 (magenta) and MAP4 (green) along microtubule tracks. The two MAPs form discrete nanoclusters along microtubule tracks. (c–e) Representative dual-colour STORM images of individual microtubule segments showing examples of MAP7D1-dominant (c), MAP4-dominant (d), and regions with both MAPs (e). (f) Pie-chart depicting the distribution of microtubule segments populated with either MAP4 or MAP7D1 or both. (g) The percentage localization density of each MAP was quantified for individual ROIs and plotted as a stacked column graph. ROIs were sorted in ascending order of MAP4 localization density, revealing an inverse correlation between MAP4 and MAP7D1 localization distribution along microtubules. (h) Representative two-colour STORM images of MAP4 and MAP7D1 nanoclusters classified into four categories based on the extent of spatial overlap. Corresponding histograms show the percentage overlap between MAP4 and MAP7D1 localizations plotted along the x-axis. (i) Pie chart shows the proportion of MAP4–MAP7D1 nanocluster overlap: non-overlapping, small overlap (<20%), partial overlap (20–80%), and high or complete overlap (>80%). This data further supports the largely non-overlapping nature of the two MAPs. Scale bars: 10 μm for (a), 5 μm for zoomed images; 1 μm for (b); 500 nm for (c–e); 100 nm for (g).

Super-resolution STORM imaging further resolved the spatial organization of MAPs, revealing that both MAP4 and MAP7D1 form discrete nanoclusters along the microtubule lattice (Figure 1b-e). Individual microtubule strands were enriched with one of the MAPs or contained both. However, the nanoclusters of MAP4 and MAP7D1 on the same microtubule exhibited minimal spatial overlap (Figure 1c–e).

To quantify their distribution, we segmented MAP-decorated microtubules into equally sized rectangular regions of interest (ROIs) (Figure 1c–e) and calculated the localization density of MAP4 and MAP7D1 within each segment. Both MAPs were imaged using Alexa Fluor 647 through sequential immunostaining and STORM imaging (see Methods), minimizing dye-dependent localization variabilities. For each segment, we computed the relative abundance of MAP4 and MAP7D1 by dividing their respective localizations by the total localizations in that segment. We defined segments with >70% MAP4 or <25% MAP7D1 as MAP4-enriched, >70% MAP7D1 or <25% MAP4 as MAP7D1-enriched, and all others as mixed. Based on this classification, ∼21% of microtubules were MAP4-enriched, ∼25% were MAP7D1-enriched, and ∼54% microtubules were co-decorated with both the MAPs (Figure 1c-f). For all the segments analyzed, notably, MAP4 and MAP7D1 densities were negatively correlated (Figure 1g), suggesting that enrichment of one MAP is correlated with depletion of the other, implying their mutually exclusive binding patterns.

We next applied HDBSCAN-based cluster analysis to determine the extent of overlap between MAP4 and MAP7D1 nanoclusters (see Methods). Each MAP4–MAP7 cluster pair was classified into one of four categories based on the percentage of spatial overlap between their localizations: (i) non-overlapping (no contact), (ii) small overlap (<20%), (iii) partial overlap (20–80%), and (iv) high or complete overlap (>80%) (Figure 1h). Among 1,024 analyzed ROIs, ∼66% of cluster pairs were largely non-overlapping, with ∼21% showing no contact and ∼45% exhibiting small overlap (Figure 1h), demonstrating a largely segregated distribution of the two MAPs. Notably, ∼32% of MAP clusters showed partial overlap, while only ∼2% displayed high or complete overlap. This spatial overlap between MAPs, although a small fraction, suggests that direct competition for microtubule binding is unlikely to account for their spatial partitioning. Interestingly, the three MAP-defined microtubule populations—enriched in either MAP4, MAP7D1, or a combination of both—mirror the distribution of microtubule PTM subpopulations quantified in our previous study^27^. We had identified three major PTM-defined subsets: ∼30% of microtubules are tyrosinated and non-acetylated, ∼35% are acetylated and tyrosinated, and ∼35% are acetylated and detyrosinated^27^. These parallel prompts the hypothesis that the distinct MAP occupancy patterns arise from differential affinities of MAP4 and MAP7D1 for specific microtubule PTM subsets.

### MAP4 prefers tyrosinated microtubules, while MAP7 is enriched on detyrosinated subset

Next, we quantitatively mapped the distribution of MAP4 and MAP7 across distinct microtubule PTM subsets marked by tyrosination, detyrosination, or acetylation in epithelial cells (BS-C-1 and HeLa) using super-resolution SIM and STORM imaging. MAP4 shows a striking preference for tyrosinated microtubules, mirroring their linear architecture while largely excluding the curved detyrosinated microtubules (Figure 2a-b; S2a-b). MAP4 binds only to a subset of acetylated microtubules (Figure 2c). Since, acetylated microtubules in BSC1 cells comprise two distinct populations — acetylated-tyrosinated and acetylated-detyrosinated microtubules^27^, we asked whether MAP4 preferentially associates with one of these subsets. Three-color SIM imaging revealed that MAP4 predominantly decorates the acetylated-tyrosinated subset and avoids acetylated-detyrosinated microtubules (Figure 2d), suggesting that acetylation does not deter MAP4’s preference for tyrosinated microtubules. Quantitative analysis confirmed this preference, with ∼90.6% of microtubule-bound MAP4 localized to tyrosinated microtubules and only ∼9.5% on detyrosinated microtubules for the same amount of microtubule PTMs (Figure 2e-f). Although acetylation does not determine MAP4’s preference between tyrosinated and detyrosinated microtubules, the concentration of MAP4 on acetylated-tyrosinated is ∼ 2folds higher compared to non–acetylated–tyrosinated microtubules, likely due to a larger proportion of acetylated-tyrosinated than non-acetylated tyrosinated microtubules. STORM imaging further revealed that MAP4 cluster density was markedly higher on tyrosinated microtubules, with sparse, scattered clusters on detyrosinated microtubules (Figure 2g-i). These results together establish that MAP4 preferentially binds to tyrosinated microtubules (Figure. 2j).

**Figure 2.**
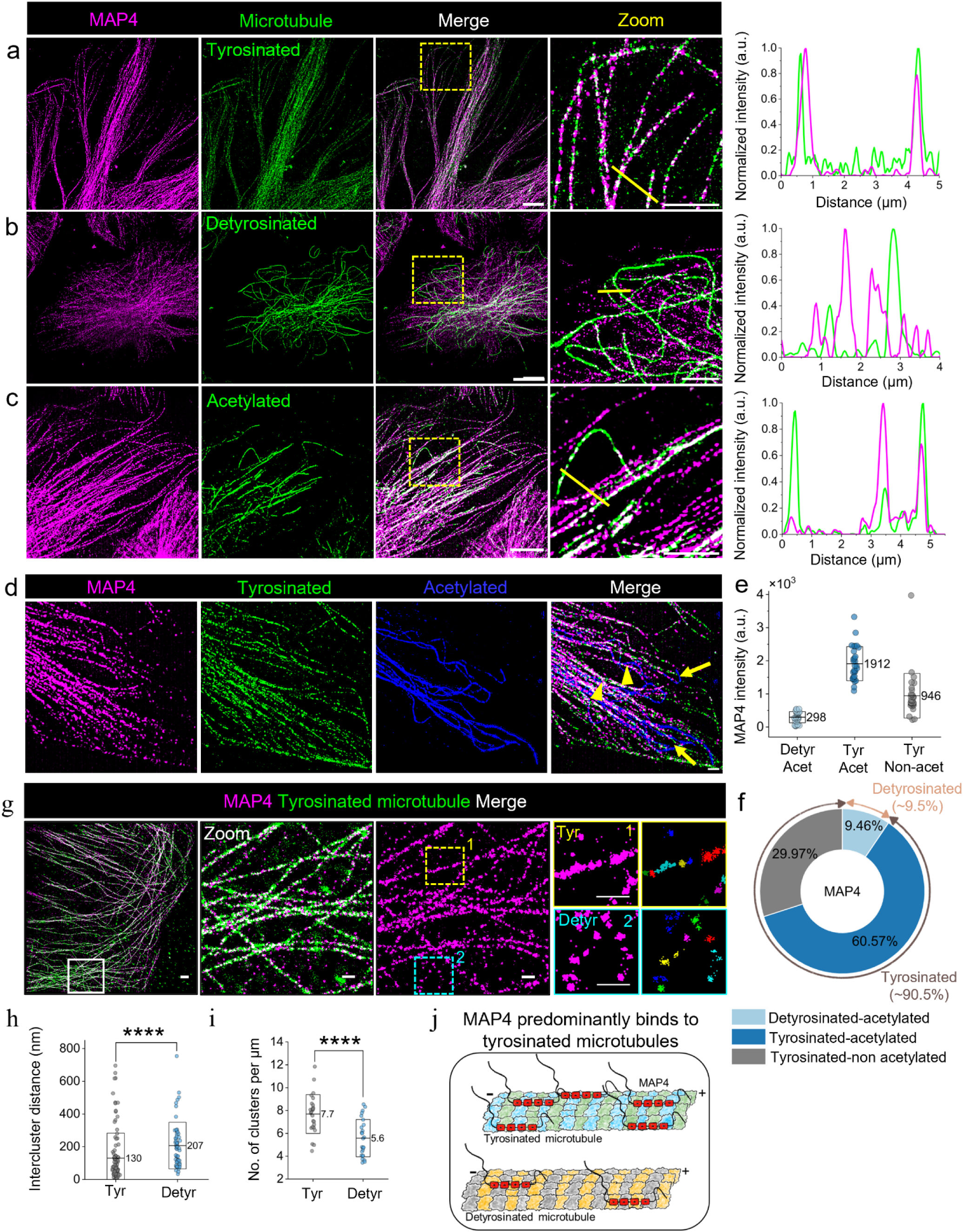
MAP4 preferentially associates with tyrosinated microtubules. Two-color super-resolution SIM images of endogenous MAP4 (magenta) with (a) tyrosinated, (b) detyrosinated, and (c) acetylated microtubules (green) in BS-C-1 cells, with corresponding line intensity plots showing preferential MAP4 localization to tyrosinated microtubules. (d) Three-color SIM image of MAP4 (magenta) with tyrosinated (green) and acetylated (blue) microtubules. Arrowheads indicate MAP4 associated with acetylated-tyrosinated microtubules; arrows indicate MAP4 associated with acetylated-detyrosinated microtubules. (e) Quantification of MAP4 localization across distinct microtubule subsets from *n* = 29 ROIs per subset from 12 different cells. MAP4 fluorescence intensity was measured from ROIs of equal size drawn on each microtubule population, ensuring uniform sampling across microtubule populations. (f) Percentage of MAP4 associated with each subset was calculated by dividing the mean MAP4 intensity on that subset by the total MAP4 intensity across all ROIs. (g) Two-color STORM images of MAP4 (magenta) with tyrosinated microtubules (green). HDBSCAN-based cluster analysis quantifies (h) inter-cluster distance and (i) cluster density (number of clusters per unit length), revealing significantly higher MAP4 clustering on tyrosinated microtubules (yellow box) compared to detyrosinated microtubules (cyan box). ROIs for detyrosinated microtubules were selected based on MAP4-decorated microtubule patterns lacking tyrosination signal. *n* = 26 ROIs were analyzed per microtubule subset from a cell. Data represents mean (line) ± SD (box). Statistical significance was assessed using the Mann–Whitney U test (****p < 0.0001). Scale bars: 10 μm for (a–c) and corresponding zoomed images at 5 μm; 5 μm for (d), 2 μm for (g), and corresponding zoomed images at 500 nm.

In contrast to MAP4, MAP7 exhibited a broader association across the subsets. In HeLa cells, endogenous MAP7 is localized to both tyrosinated and detyrosinated microtubules, although with a striking similarity with the architecture of detyrosinated microtubules (Figure S3a). Similarly, in BSC-1 cells, the predominant microtubule-bound isoform, MAP7D1 (Figure S1a), is distributed across both tyrosinated and acetylated microtubules in a differential manner (Figure 3a-b). Although antibody cross-reactivity precluded direct visualization of MAP7D1 with detyrosinated microtubules, three-color SIM imaging with acetylation and tyrosination markers enabled us to infer acetylated-detyrosinated regions (Figure 3c) and quantify the distribution of MAP7D1 on these PTM subsets. Notably, ∼58.4% of MAP7D1 localized to acetylated-detyrosinated microtubules, ∼32.4% to acetylated-tyrosinated microtubules, and only ∼9.2% to tyrosinated microtubules lacking acetylation (Figure 3d-e), suggesting that MAPD1 has an affinity for detyrosinated microtubules and further acetylation likely enhances MAP7D1 binding with tyrosinated microtubules. Complementary STORM imaging revealed a distinct nanoscale organization of MAP7 isoform overexpressed in BS-C-1 cells, with dense MAP7 nanoclusters on detyrosinated microtubules and sparsely distributed clusters on tyrosinated microtubules (Figure 3f-h).

**Figure 3.**
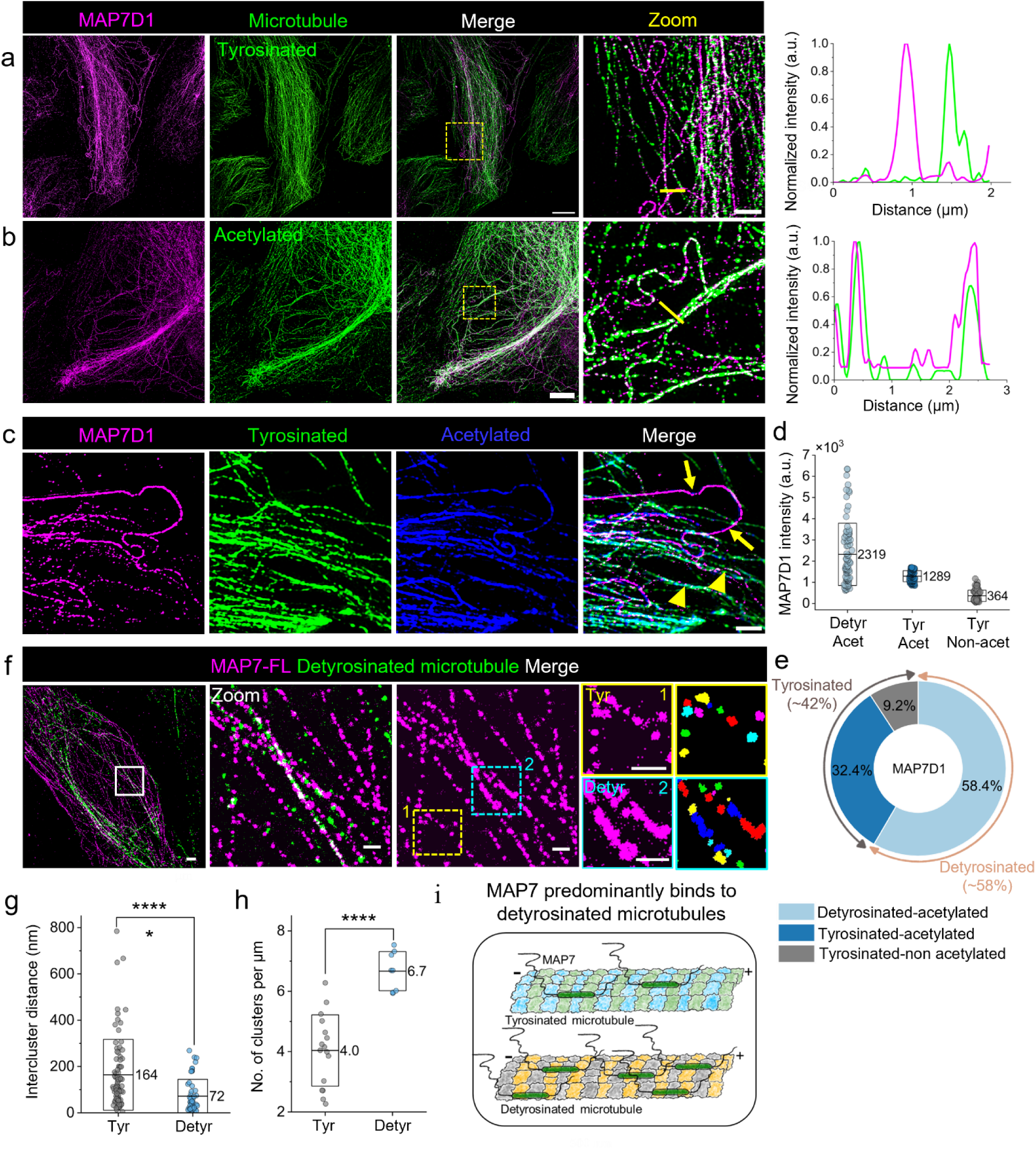
MAP7D1 is enriched on detyrosinated microtubules. Two-color SIM images of endogenous MAP7D1 (magenta) with (a) tyrosinated and (b) acetylated microtubules (green) in BS-C-1 cells, with corresponding line intensity plots showing MAP7D1 localization to both the subsets. (c) Three-color SIM image of endogenous MAP7D1 (magenta) with tyrosinated (green) and acetylated (blue) microtubules. Arrows indicate MAP7D1 enriched on acetylated-detyrosinated microtubules, and arrowheads indicate MAP7D1 associated with acetylated-tyrosinated microtubules. (d) Quantification of MAP7D1 localization across distinct microtubule subsets from n=82 ROIs per subset from 11 different cells. MAP7D1 fluorescence intensity was measured from ROIs of equal size drawn on each microtubule population, ensuring uniform sampling across microtubule populations. (e) The percentage of MAP7D1 associated with each subset was calculated by dividing the mean MAP7D1 intensity on that subset by the total MAP7D1 intensity across all ROIs. (f) Two-color STORM images of overexpressed full-length MAP7 (magenta) and detyrosinated microtubules (green) in BS-C-1 cells. HDBSCAN-based cluster analysis quantifies (g) inter-cluster distance and (h) cluster density (number of clusters per unit length), revealing significantly higher MAP7 clustering on detyrosinated microtubules (cyan box) compared to tyrosinated microtubules (yellow box). ROIs for tyrosinated microtubules were selected based on MAP7-decorated microtubule patterns lacking detyrosination signal. *n* = 16, and 8 ROIs were analyzed for tyrosinated and detyrosinated microtubule subsets. Data represents mean (line) ± SD (box). Statistical significance was assessed using the Mann–Whitney U test (****p < 0.0001). Scale bars: 10 μm for (a, b), with zoomed images at 5 μm; 5 μm for (d), 2 μm for (f), with zoomed images at 500 nm.

Together, these findings reveal the distinct and contrasting microtubule localization of MAP4 and MAP7. MAP4 is preferentially associated with tyrosinated microtubules, while MAP7 is enriched on detyrosinated microtubules (Figure 2i, 3j). This specific pattern of MAP distribution on microtubules suggests a regulatory crosstalk between tubulin-modifying enzymes and microtubule-bound MAPs.

### MAP7D1 prefers expanded lattices, while MAP4 selects tyrosinated microtubules independent of lattice state

We investigated whether tubulin modifications act as spatial cues for distinct MAP-microtubule associations. Tubulin tyrosine ligase (TTL) and vasohibin (VASH1/2)–SVBP complex regulate the cellular balance of tyrosinated and detyrosinated microtubules^41,42^. Overexpression of VASH2 robustly increased detyrosinated microtubules (Figure S4a), driving widespread MAP4 localization onto them (Figure S4b). This indicates that MAP4’s preference for tyrosinated microtubules is not due to detyrosination preventing its binding. Conversely, reducing the overall detyrosinated microtubules by increasing tyrosination through TTL overexpression (Figure S4a) led MAP7D1 to bind to tyrosinated microtubules, however it did not equally distribute MAP7D1 on all microtubules, rather maintained its enrichment on a subset of microtubules (Figure S4c), suggesting that detyrosination alone is not likely to be the determinant for MAP7D1 enrichment on a small subset of microtubules.

This raises the possibility of coordinated mechanisms that regulate both microtubule-PTMs and MAP binding. Given that the microtubule lattice conformation, expanded or compacted, can influence the binding of both MAPs and PTM enzymes ^37,38,43,44^, we asked whether the microtubule lattice state contributes to MAP-PTM specificity. To test this, we induced lattice expansion by treating BS-C-1 cells with Taxol (100 µM, 10 min) and examined MAP localization relative to DMSO controls (Figure 4a-d). Notably, MAP4 retained its strong preference for tyrosinated microtubules upon lattice expansion (Figure 4a, b). In contrast, MAP7D1, typically enriched on detyrosinated microtubules, is redistributed across all microtubule subsets, including the tyrosinated ones (Figure 4c, d). Next, we performed STORM imaging of MAP4 and MAP7D1 after Taxol or DMSO treatment (Figure 4e) and analysed their distribution on tyrosinated microtubules as identified from the corresponding two-colour TIRF imaging of MAP4 or MAP7D1 with tyrosinated microtubules (Figure S4d, e). Quantitative analysis confirmed that MAP4 localization density remained unaltered, whereas MAP7D1 significantly increased its density on tyrosinated microtubules following expansion (Figure 4e, f). These findings indicate that MAP7D1 preferentially associates with expanded lattice, a conformation often coupled with detyrosination, consistent with prior reports of VASH1’s affinity for expanded microtubules^38^.Together, our results reveal distinct modes of selective MAP-PTM association. MAP7D1 is enriched on the expanded microtubule lattice, which also accumulates detyrosination. In contrast, MAP4’s selective association with tyrosinated microtubules is not determined by lattice conformation and detyrosination status. This suggests that MAP4 might harbor intrinsic structural features that either promote its binding to tyrosinated microtubules or act to preserve microtubule tyrosination by restricting detyrosination at its binding sites.

**Figure 4:**
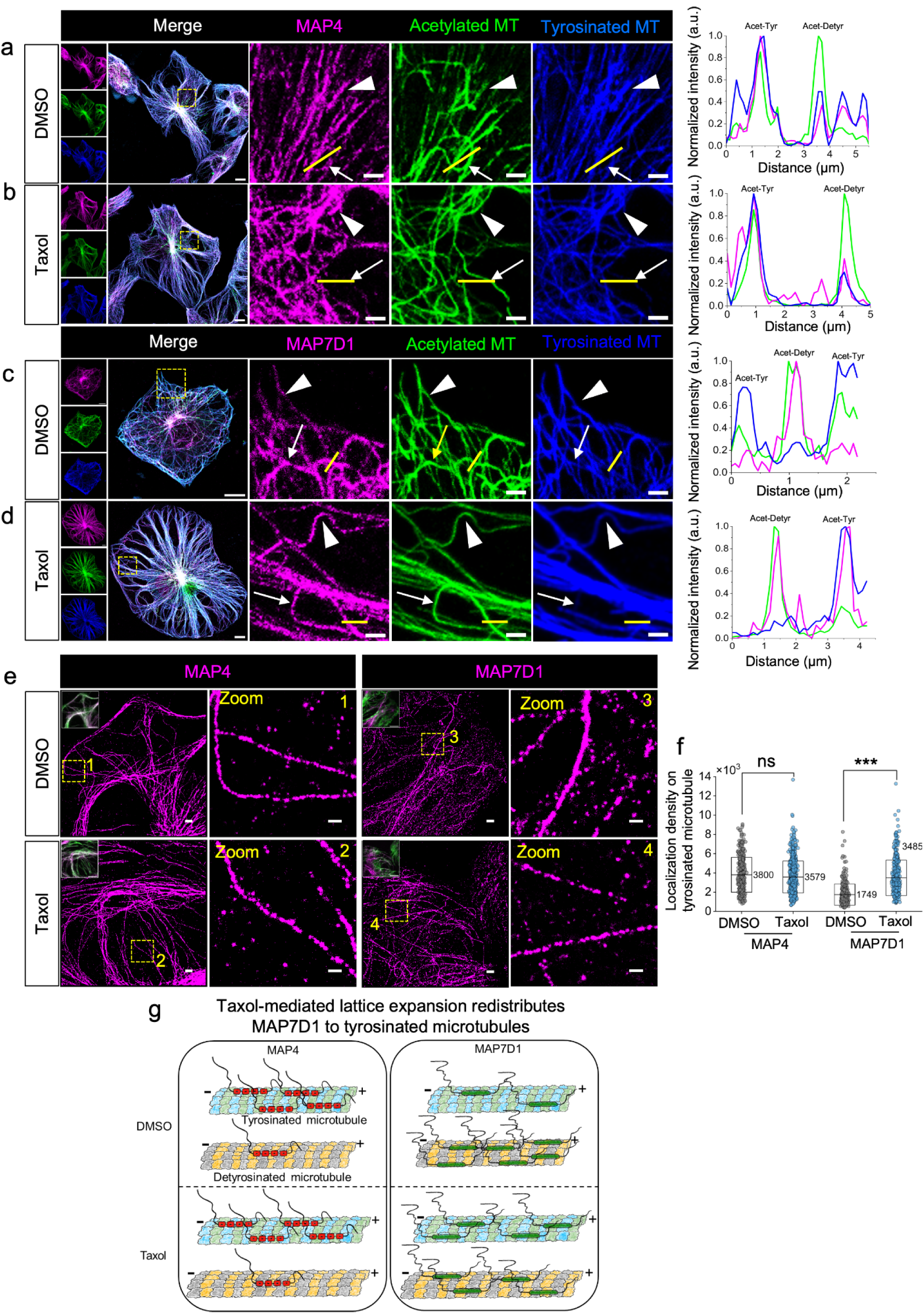
MAP7D1, not MAP4, is sensitive to lattice expansion. (a-d) Three-color confocal images and corresponding line intensity profiles of BS-C-1 cells (a) DMSO-treated cells immunostained for endogenous MAP4 (magenta) with acetylated (green) and tyrosinated (blue) microtubules. Arrowheads indicate MAP4 colocalization with acetylated-tyrosinated microtubules; arrows indicate MAP4 exclusion from acetylated-detyrosinated microtubules. (b) Taxol-induced lattice expansion preserves MAP4’s preference for acetylated-tyrosinated microtubules. (c) DMSO-treated cells immunostained for endogenous MAPD1 (magenta) with acetylated (green) and tyrosinated (blue) microtubules. Arrowheads indicate MAP7D1 sparsely decorated on acetylated-tyrosinated microtubules; arrows indicate MAP7D1 enriched on acetylated-detyrosinated microtubules. (d) Taxol-induced lattice expansion redistributes MAP7D1 to acetylated-detyrosinated (arrows) and acetylated-tyrosinated (arrowheads) microtubules. (e) Super-resolution STORM images showing endogenous MAP4 (magenta) or MAP7D1 (magenta) localization on tyrosinated microtubules in DMSO or Taxol-treated cells. ROIs were drawn on tyrosinated microtubules based on corresponding two-color TIRF images (shown as insets, also see Figure S3b, c). Yellow box shows the corresponding zoomed ROIs, highlighting unaltered MAP4 localization but increased MAP7D1 density on tyrosinated microtubules after lattice expansion. (h) Quantification of MAP4 and MAP7D1 localization density on tyrosinated microtubules from STORM images. No. of ROIs analyzed, MAP4 DMSO (*n*=383), MAP4 Taxol (n= 658), MAP7D1 DMSO (n= 317), MAP7D1 Taxol (n= 374) from 3 different cells. Data represents mean (line) ± SD (box). Statistical significance was assessed using the Mann–Whitney U test (****p < 0.0001, not significant [ns]). Scale bars: 10 μm for (a–d), with zoomed images at 2 μm; 2 μm for (e), with zoomed images at 500 nm.

### MAP4 projection domains define its selectivity for tyrosinated microtubules

To identify MAP4’s structural determinants that confer its preference for tyrosinated microtubules, we generated a series of GFP-tagged truncation variants of full-length MAP4 (MAP4-FL): MAP4-ΔC (lacking the short C-terminal projection domain), MAP4-ΔN (lacking the long N-terminal projection domain), and MAP4-MTBD (containing only the microtubule-binding domain) (Figure 5a, S5a). These constructs were transiently expressed in BS-C-1 cells to assess their localization to detyrosinated microtubules (Figure 5b–f).

**Figure 5:**
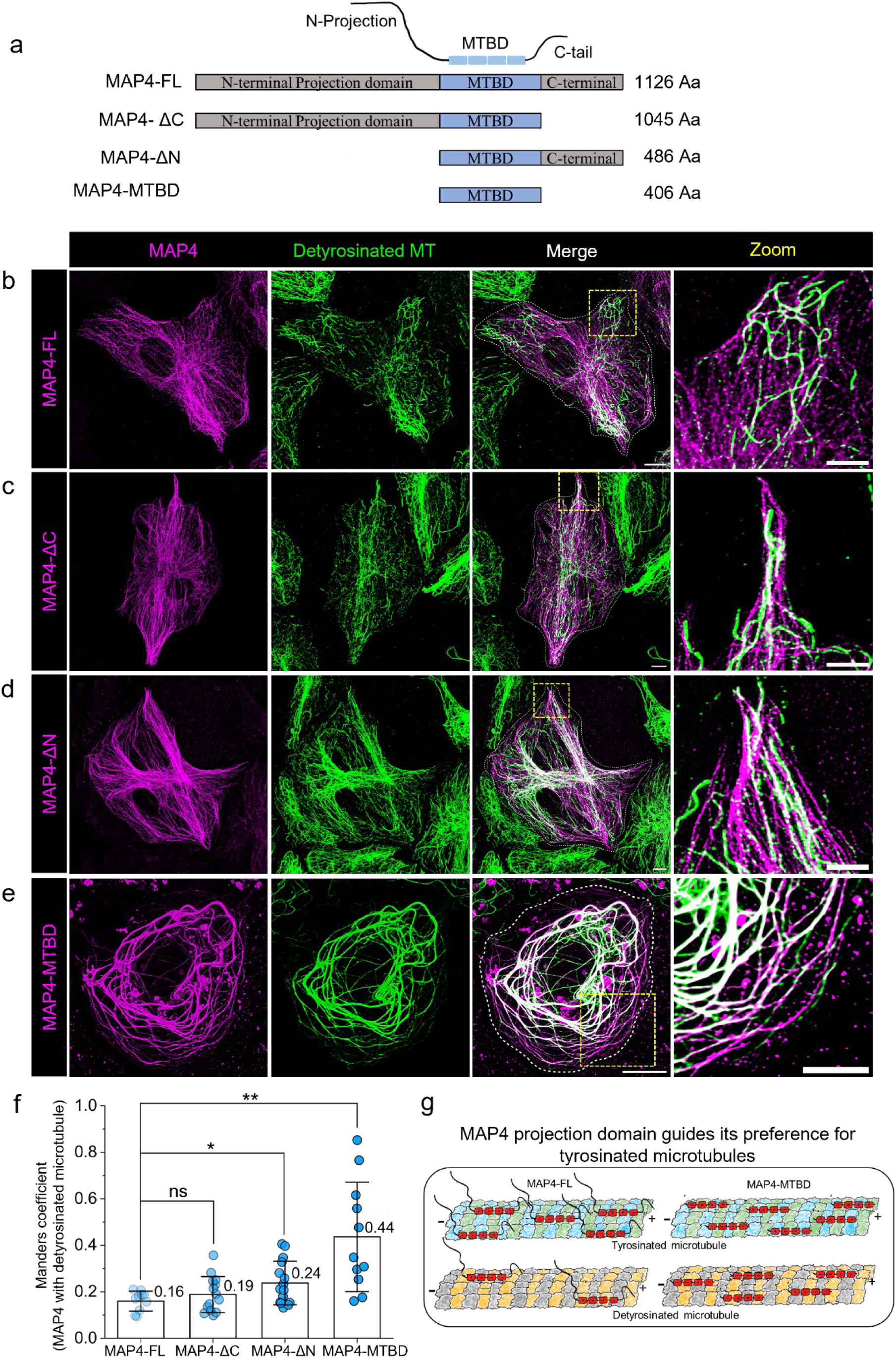
MAP4 projection domains restrict its association to tyrosinated microtubules. (a) The Schematic of MAP4 truncation constructs used in the study. (b-e) Representative confocal images of BS-C-1 cells transiently expressing GFP-tagged MAP4 constructs (magenta) immunostained for detyrosinated microtubules (green) along with corresponding insets show magnified views of the regions indicated by yellow boxes. (f) Box plot quantifying MAP4 colocalization with detyrosinated microtubules. (g) Model illustrating the role of MAP4 projection domains in conferring specificity for tyrosinated microtubules. Data represents mean (line) ± SD (box). Number of cells (n) analyzed for MAP4-FL (10), MAP4-MTBD (11), MAP4-ΔN (15) and MAP4-ΔC (14). Statistical significance was assessed using the Mann– Whitney U test (*p<0.05, **p<0.01, not significant [ns]). Scale bars: 10 μm.

Notably, overexpressed MAP4-FL remained largely excluded from detyrosinated microtubules (Figure 5b, f), indicating that elevated MAP4 levels do not override its intrinsic preference for tyrosinated microtubules. Strikingly, progressive deletion of the projection domains increased MAP4 localization on detyrosinated microtubules — with MAP4-ΔC showing modest localization, MAP4-ΔN showing greater overlap, and MAP4-MTBD exhibiting the highest colocalization (Figure 5c–e). Quantification confirmed that deletion of the N-terminal projection domain (MAP4-ΔN) significantly enhanced MAP4 association with detyrosinated microtubules compared to C-terminal deletion alone (MAP4-ΔC) (Figure 5f). These results implicate the N-terminal projection domain as a key determinant restricting MAP4 binding to tyrosinated microtubules (Figure 5g).

Additionally, the deletion of MAP4 projection domains led to an increase in cellular detyrosinated microtubules, with MAP4-MTBD expression having significantly higher detyrosination levels than MAP4-FL (Figure S5b). Collectively, these findings suggest that MAP4’s projection domain governs its selective association with tyrosinated microtubules and shields MAP4-decorated microtubules from detyrosination. Additionally, plausible interactions of the projection domain with other microtubule-associated factors present on detyrosinated microtubules prevent MAP4 from binding to pre-existing detyrosinated microtubules.

### MAP4 associates with kinesin-3 while MAP7D1 localizes with kinesin-1

MAPs regulate motor proteins majorly through their projection domains. Given that ATPase-deficient rigor mutants of kinesin-1 and kinesin-3 preferentially bind to distinct microtubule PTMs^6^, we investigated whether specific patterns of MAPs–PTMs are associated with specific motor proteins. We transiently expressed rigor mutants of KIF5B-R (kinesin-1) and KIF1A-R (kinesin-3) in BS-C-1 cells and examined their localization relative to tyrosinated and detyrosinated microtubules. Consistent with previous reports^6^, KIF5B-R preferentially decorated detyrosinated microtubules, whereas KIF1A-R localized predominantly to rectilinear tyrosinated microtubules (Figure S6a).

Strikingly, we observed a significantly high colocalization between MAP4 and KIF1A-R on tyrosinated microtubules as compared to MAP4 and KIF5B-R, which are associated with mutually exclusive microtubule subsets (Figure 6a, c). In contrast, MAP7D1 showed the opposite pattern — colocalizing robustly with KIF5B-R on detyrosinated microtubules compared to KIF1A-R (Figure 6d-f). Interestingly, KIF5B-R expression further enriched MAP7D1 localization on detyrosinated microtubules (Figure S6b), likely through motor-induced lattice expansion^45^.

**Figure 6.**
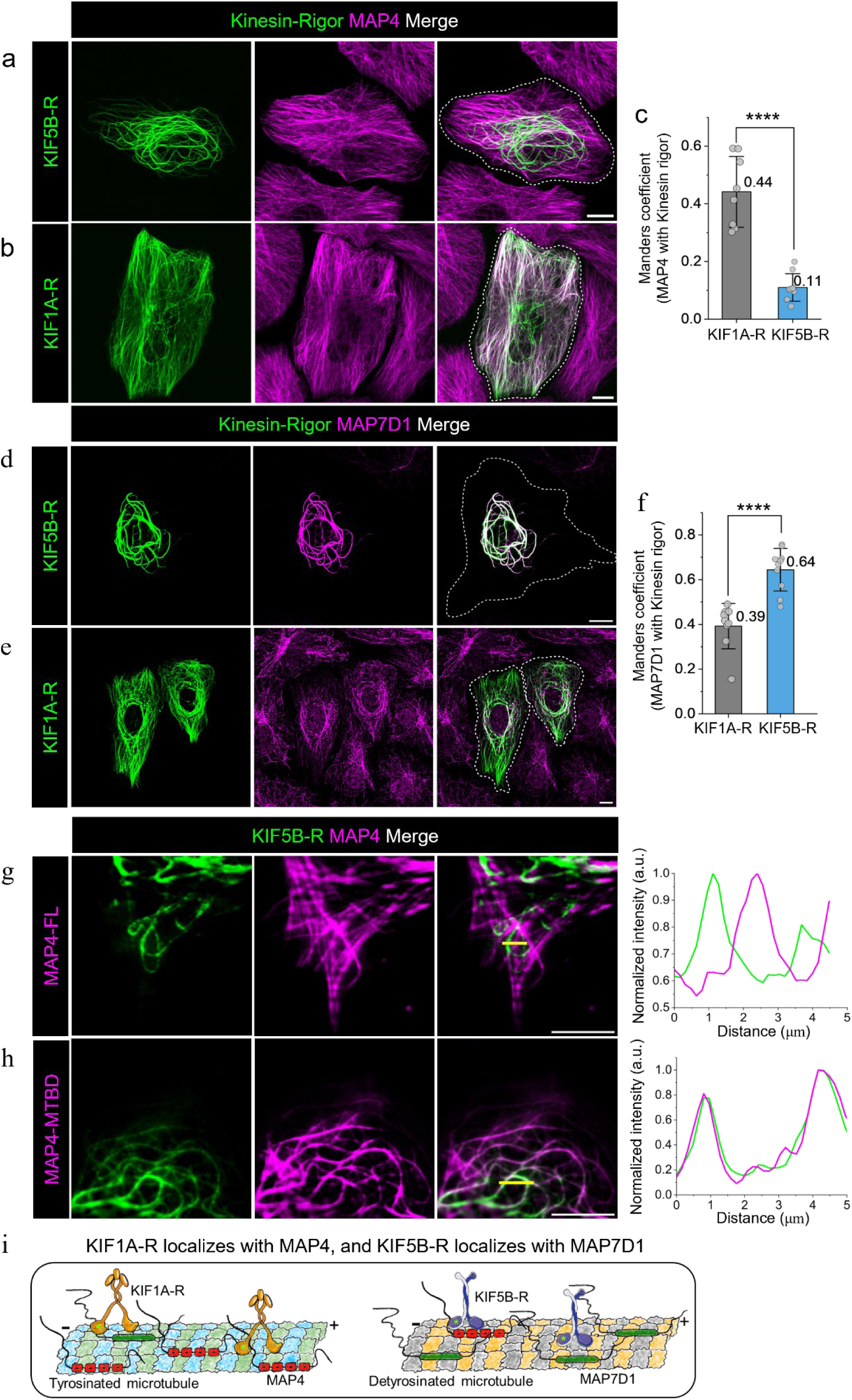
Kinesin-1 and kinesin-3 rigor mutants differentially associate with MAP4 and MAP7D1 (a-b) Confocal images of BS-C-1 cells transiently expressing GFP-tagged (a) KIF5B-R (green) or (b) KIF1A-R (green), immunostained for endogenous MAP4 (magenta). (c) Bar plot of Mander’s colocalization quantification shows significantly higher colocalization of MAP4 with KIF1A-R (*n* = 8 cells) than KIF5B-R (*n* = 9 cells). (d-e) Confocal images of cells expressing GFP-tagged (d) KIF1A-R or (e) KIF5B-R (green), immunostained for endogenous MAP7D1 (magenta). (f) Bar plot of Mander’s colocalization quantification reveals significantly higher colocalization of MAP7D1 with KIF5B-R (*n* = 10 cells) than KIF1A-R (*n* = 9 cells). (g–h) Co-expression of KIF5B-R (green) with (g) MAP4-FL or (h) MAP4-MTBD (magenta), Line intensity profiles show MAP4-FL is excluded from KIF5B-R-decorated microtubules, while MAP4-MTBD exhibits extensive colocalization. (i) Model illustrating the preferential association of KIFB-R and KIF1A-R with MAP7D1 and MAP4, respectively. Bars represent the mean; whiskers indicate standard deviation from three independent experiments. Statistical significance was assessed using the Mann–Whitney U test (****p < 0.0001). Scale bars: 10 μm.

To test whether MAP4’s projection domain governs its exclusion from KIF5B-decorated microtubules, we co-expressed KIF5B-R with either full-length MAP4 (MAP4-FL) or the truncated variant lacking both projection domains (MAP4-MTBD). Consistent with our hypothesis, MAP4-FL was excluded from KIF5B-R-decorated microtubules, whereas MAP4-MTBD showed substantial colocalization with KIF5B-R (Figure 6d-e). These findings suggest that MAP4 associates with KIF5B in a projection domain-dependent manner.

Together, these results highlight that MAP4 and MAP7D1 differentially interact with kinesin-1 and kinesin-3 to coordinate organelle transport.

### Discrete MAP nanoclusters on microtubules regulate lysosome motility in a concentration and direction-dependent manner

Super-resolution STORM images show that MAP4 and MAP7D1 form discrete nanoclusters on the microtubule lattice (Figure 1b-e, 2g, 3f). We used an all-optical correlative live-cell and STORM imaging approach to determine how these discrete MAP clusters regulate motor-mediated lysosome movement. In BS-C-1 cells stably expressing LAMP1-YFP, lysosome motility was recorded at 10 fps for ∼1 minute using live-cell TIRF imaging, followed by *in situ* immunostaining and STORM imaging of endogenous MAP4 or MAP7D1. Lysosome tracks were overlaid on STORM images of MAP4 or MAP7D1 (Figure 7a, e). TetraSpeck beads were used as fiduciary markers for image registration and alignment of live-cell TIRF and STORM imaging channels (Figure S7a).

**Figure 7.**
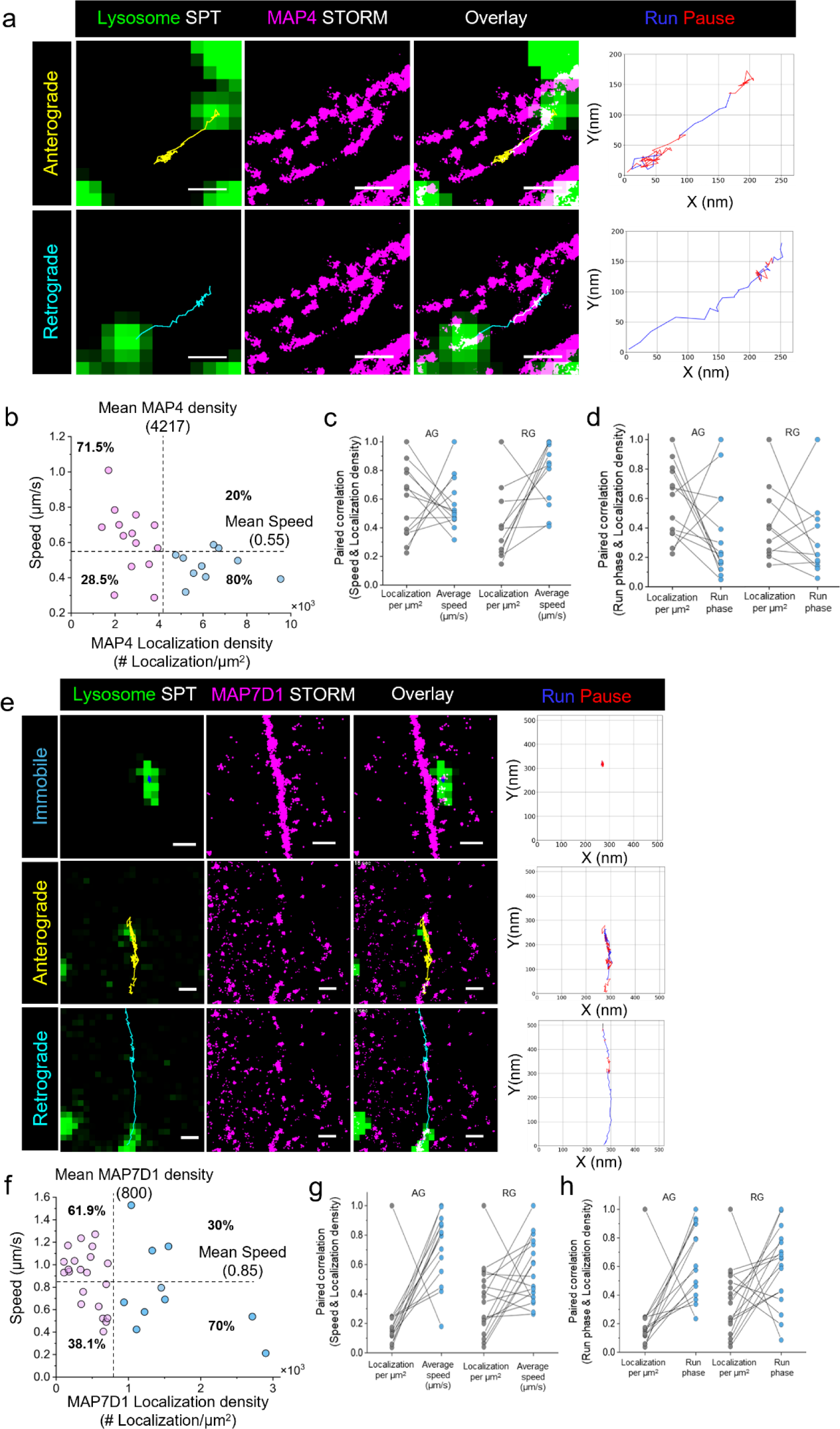
MAP4 and MAP7D1 nanoclusters regulate lysosome motility in a concentration- and direction-dependent manner. (a) Correlative live-cell and STORM imaging of lysosomes with MAP4 in BS-C-1 cells. Single-particle trajectories (SPT) from live imaging were aligned and overlaid on the corresponding STORM image of MAP4. Anterograde track is shown in yellow and retrograde track in cyan. In corresponding graphs, the trajectories are colour-coded to indicate run phases (blue) versus pause phases (red), revealing more processive retrograde runs than anterograde runs. (b) Scatter plot of lysosome speed versus MAP4 density. Dotted lines mark the mean on both axes. Points below (pink) or above (blue) the mean MAP4 density illustrate that lysosomes show reduced average speed at higher MAP4 density. (c) Paired correlation plot between lysosomal speed and MAP4 density, separated into anterograde (n = 13 trajectories) and retrograde (n = 11 trajectories) tracks. This shows that anterograde-moving lysosomes experience greater hindrance in speed with increasing MAP4 density. (d) Paired correlation plot between run-phase and MAP4 density for the same anterograde (n = 13) and retrograde (n = 11) tracks. Higher MAP4 density reduces run-phase more markedly for anterograde than for retrograde tracks. (e) Correlative live-cell and STORM imaging of lysosomes with MAP7D1 (magenta) in BS-C-1 cells. Single-particle trajectories of lysosomes from live imaging were aligned and overlaid on the STORM image of MAP7D1. Anterograde trajectories are marked in yellow and retrograde in cyan. Run phases (blue) and pause phases (red) highlight that retrograde runs are more processive than anterograde runs. (f) Scatter plot of lysosome speed versus MAP7D1 density. Dotted lines mark the mean on both axes. Points below (pink) or above (blue) the mean MAP7D1 density indicate that lysosomes have reduced average speed at higher MAP7D1 density. (g) Paired correlation plot between lysosomal speed and MAP7D1 density, separated into anterograde (n = 14 trajectories) and retrograde (n = 17 trajectories) tracks. Anterograde-moving lysosomes show greater hindrance in speed as MAP7D1 density increases. (h) Paired correlation plot between run-phase and MAP7D1 density for the same anterograde (n = 14) and retrograde (n = 17) tracks. Higher MAP7D1 density reduces run-phase measures more for anterograde than retrograde tracks. Scale bars: 500 nm.

We classified the lysosome trajectories as anterograde or retrograde based on the track direction towards or away from the nucleus. Using a custom-written algorithm (see methods for more details), we segmented lysosome trajectories into processive (run) and non-processive (pause) phases. We observe that as the lysosome moves on a MAP4 decorated microtubule, it exhibits more non-processive pause behaviour in the anterograde direction, as compared to the lysosomes moving in the retrograde direction, for the same localization density of MAP4 (Figure 7a, Movie S2, S3), suggesting that higher concentrations of MAP4 exert more inhibition on kinesin-driven lysosome motility than dynein. Next, we quantified the speed of the lysosomal trajectories and correlated them with the respective MAP4 localization density along each trajectory. Interestingly, below the mean MAP4 density (4.2×10^3^ localizations/μm²), 71% of lysosomes moved faster than the average speed (0.55 μm/s), suggesting that MAP4 favours lysosomal transport at optimal densities. In contrast, with above-average MAP4 densities, only 20% of lysosomes exceeded the average speed, indicating that high MAP4 density impedes lysosome movement (Figure 7b). Further directional analysis revealed that the speed and the run-phase of the lysosomes moving in anterograde (AG) direction were more impeded by high MAP4 density as compared to retrograde (RG) movement (Figure 7c, d, Movie S2, S3). Furthermore, increasing the cellular MAP4 concentration by MAP4-FL overexpression resulted in decreased lysosomal motility and increased proportion of immobile lysosomes (Figure S7b-d). Thus, MAP4 regulate lysosome motility in a concentration and direction-dependent manner.

Similarly, we observe that lysosomes are largely immobile on microtubules decorated with very high MAP7D1 density (Figure 7e). For the same local density of MAP7D1, lysosomes exhibited more impaired AG than RG motility (Figure 7e, Movie S5, S6). Below the mean density of MAP7D1 (0.8×10^3^ localizations/μm^2^), ∼62% of lysosomes moved with speed faster than average speed (0.85 μm/s), whereas only ∼30% of lysosomes moved with average speed beyond the mean MAP7D1 density (Figure 7f). Akin to MAP4, high MAPD1 density exhibited stronger impedance to AG than RG motility of lysosomes (Figure 7g, h), and overexpression of MAP7 FL resulted in overall impairment of lysosome motility and an increase in immobile lysosome population (Figure S7b-d). Our findings, thereby, suggest that MAP4 and MAP7 densities fine-tune the motility of lysosomes and control their directionality. Additionally, we observed intriguing lysosomal trafficking behaviors upon encountering MAP4 or MAP7D1 clusters. At the MAP4 and MAP7D1 interception points, lysosomes either paused before resuming movement (Figure S7e, Movies S2, S6, S7) or passed through without interruption (Figure S7f, Movies S3, S8, S9). Following a pause, lysosomes display diverse bypass strategies, including hopping to the next MAP cluster (Figure S7e, Movie S7, S8), or switching to an adjacent microtubule track (Figure S7f, Movie S9, S10), to circumvent the MAP obstacle. Notably, we did not observe lysosome track reversals upon MAP encounter. Although hopping and track switching events were less frequent, likely because endogenous MAP concentrations are permissive for motor passage rather than acting as roadblocks, these distinct trafficking behaviors suggest that at sub-optimal MAP densities, lysosomes may employ such motor-specific mechanisms to overcome MAP obstacles on microtubules. Future investigations with specifically defined MAP–motor interactions is critical to uncover the mechanistic basis of how lysosomes overcome MAP-mediated roadblocks during intracellular transport.

### MAP density on microtubules regulates nutrient-dependent remodelling of lysosomes

Lysosomal motility is differentially regulated by the density of MAPs on microtubules. However, whether cells actively tune MAP density to modulate lysosomal trafficking and cytoplasmic distribution has remained unclear. Previous studies have demonstrated that lysosomes undergo dynamic repositioning in response to nutrient availability, a process critical for lysosomal function and driven by long-range motor-dependent transport along microtubules.

Consistent with previous observations, nutrient starvation led to a pronounced accumulation of lysosomes in the perinuclear region, whereas stimulation with 2× amino acids for 20 minutes triggered rapid redistribution of lysosomes to the cell periphery compared to untreated control cells (Figure 8a, b). Beyond repositioning, nutrient starvation markedly increased lysosomal size while reducing their number, whereas nutrient stimulation increased lysosome number and decreased their size, most likely due to lysosomal fission (Figure S8a–c). These results establish that lysosomes undergo rapid remodelling in response to nutrient fluctuations.

**Figure 8:**
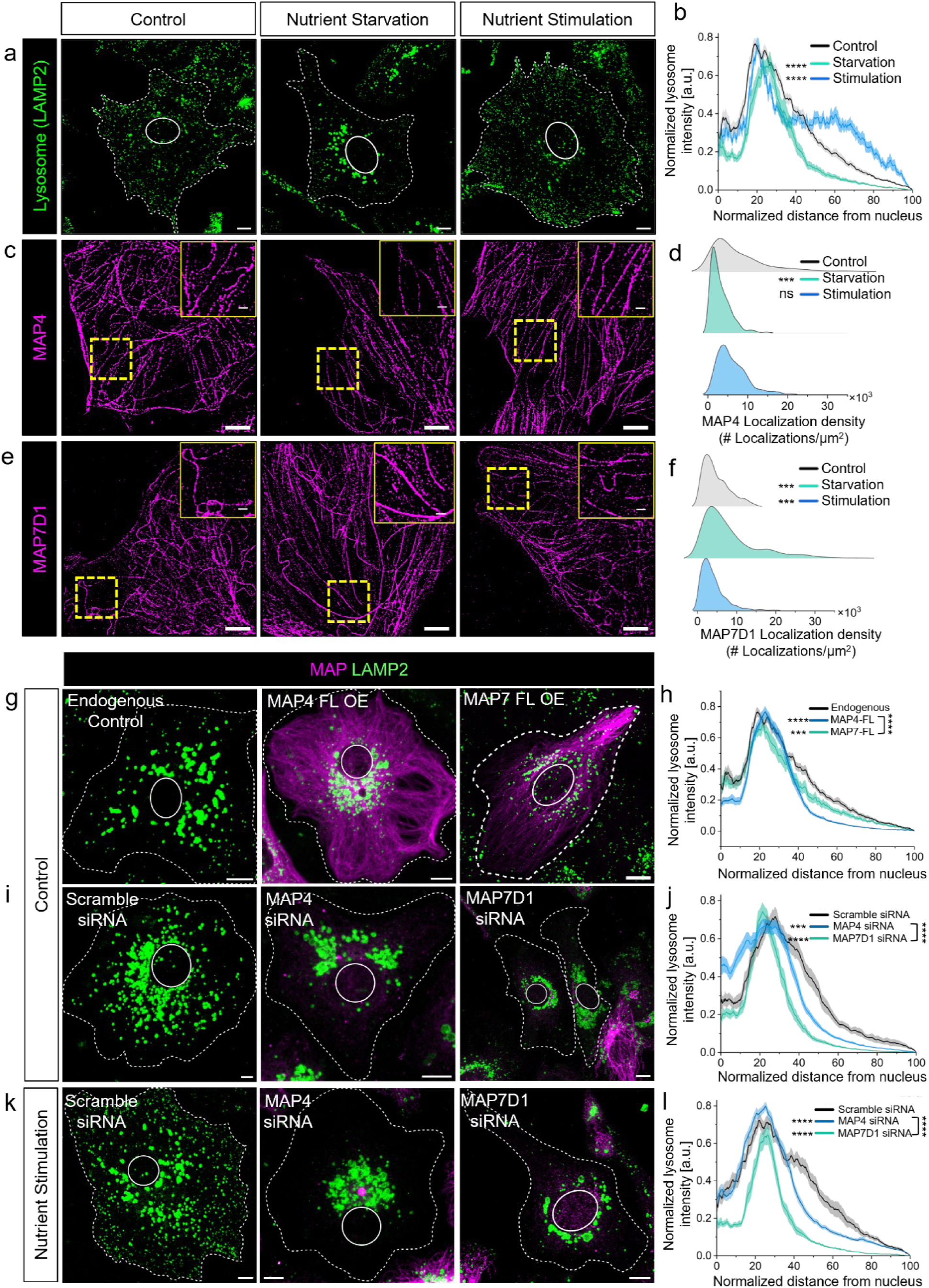
MAP density on microtubules regulates nutrient-dependent lysosomal positioning (a) Representative confocal images of BS-C-1 cells immunostained for endogenous LAMP2 (green) showing lysosomal distribution and (b) radial intensity profile plot for quantification of lysosomal distribution under control (n=27 cells), nutrient starvation (n=17 cells) and 2x amino acids stimulated (n=17 cells) conditions, (c-f) Representative STORM images MAP4 and MAP7D1 under control, nutrient starvation and nutrient stimulation along with corresponding localization density distribution plots from 3 different cells in each conditions. (g-h) Representative confocal images of control cells showing lysosomal distribution along with the corresponding radial intensity profile plot in endogenous (n=27 cells), MAP4-FL overexpression (n=27 cells) and MAP7-FL overexpression (n=23 cells) conditions. (i-j) Representative confocal images of untreated control cells showing lysosomal distribution along with corresponding radial intensity profile plot in scramble siRNA (n=21 cells), MAP4 siRNA (n=37 cells) and MAP7D1 siRNA (n=27 cells). (k-l) Representative confocal images of nutrient-stimulated cells showing lysosomal distribution along with corresponding radial intensity profile plot in scramble siRNA (n=23 cells), MAP4 siRNA (n=41 cells) and MAP7D1 siRNA (n=31 cells). Lysosomal distribution as a function of distance from the center of the nucleus was quantified from confocal images using the Radial Profile plugin in ImageJ (see Methods for details). Radial intensity profile graphs represent the mean ± SEM of lysosomal intensity profiles, averaged from three independent experiments. Statistical significance was assessed using the Kolmogorov–Smirnov (KS) test, comparing treated conditions to the control (n.s.> 0.05, *** < 0.05, **** p < 0.00001). Scale bars: 10 μm for confocal images, 5 μm for STORM images in c, e and corresponding insets at 1 μm.

Since MAP concentration locally regulates the directionality of motor-mediated transport, we next asked whether nutrient-driven lysosomal remodelling correlates with changes in MAP density. Dual-colour confocal imaging of MAP4 and MAP7D1 in BS-C-1 cells under control, starvation, and nutrient stimulation conditions revealed striking differences in their distribution (Figure S8d-e). Quantitative super-resolution STORM imaging further showed that nutrient starvation significantly decreased MAP4 localization density on microtubules while increasing MAP7D1 density (Figure 8c, d, S8d, f). Conversely, nutrient stimulation reduced MAP7D1 density but left MAP4 density largely unaltered (Figure 8e, f, S8e, g). These results suggest cellular mechanisms that dynamically tune MAP concentration on microtubules in response to nutrient state, which likely facilitates lysosomal positioning accordingly.

To directly test whether MAP modulation is required for lysosomal reorganization, we perturbed MAP levels in cells Transient overexpression of full-length MAP4 or MAP7 led to a significant perinuclear accumulation of lysosomes, with MAP4 overexpression exhibiting a more pronounced phenotype (Figure 8g, h, S8g). This reflects our observations that at higher concentrations, MAP4 is more inhibitory for lysosomal anterograde transport than for retrograde transport. Conversely, siRNA-mediated knockdown of MAP4 or MAP7D1 also induced perinuclear clustering of lysosomes relative to scramble controls, with MAP7D1 depletion resulting in more substantial lysosome accumulation near the nucleus (Figure 8i, j; Figure S8g). Previous studies have shown that MAP7 is essential for kinesin-1 recruitment and activation to steer cargo towards cell periphery, partly explaining the perinuclear accumulation of lysosomes we observe with MAP7D1 knockdown. Distinct and possibly more complex mechanisms likely underlie the interaction between specific kinesins and dynein with MAP4 and MAP7, regulating lysosomal positioning, which warrants further investigation of these molecular interactions using isolated protein systems. Nonetheless, our results indicate that an optimal density of MAP4 and MAP7 on microtubules is required to fine-tune the balance between anterograde and retrograde transport and positioning of lysosomes. Strikingly, nutrient stimulation in cells depleted of MAP4 or MAP7D1 failed to promote lysosomal redistribution to the periphery (Figure 8k, l, S8g). This demonstrates that optimal MAP density on microtubules is critical for nutrient-responsive lysosomal repositioning. Together, these findings demonstrate that cells dynamically regulate MAP density on microtubules in response to fluctuations in nutrient availability. This mechanism ensures that lysosomal positioning and remodelling remain permissive to motor-driven transport under changing metabolic conditions.

## Discussion

Our study uncovers a spatial logic by which distinct microtubule post-translational modifications—tyrosination, detyrosination, and acetylation segregate MAP4 and MAP7, establishing functional MAP gradients that differentially regulate kinesin-1 and kinesin-3 motors. This molecular compartmentalization coordinates lysosomal positioning and transport, integrating microtubule identity with nutrient-responsive signalling pathways (Figure 9). These specific MAP–PTM combinations emerge from a complex interplay between MAP–PTM enzyme interactions and enzyme accessibility along the structural state of the microtubule lattice.

**Figure 9.**
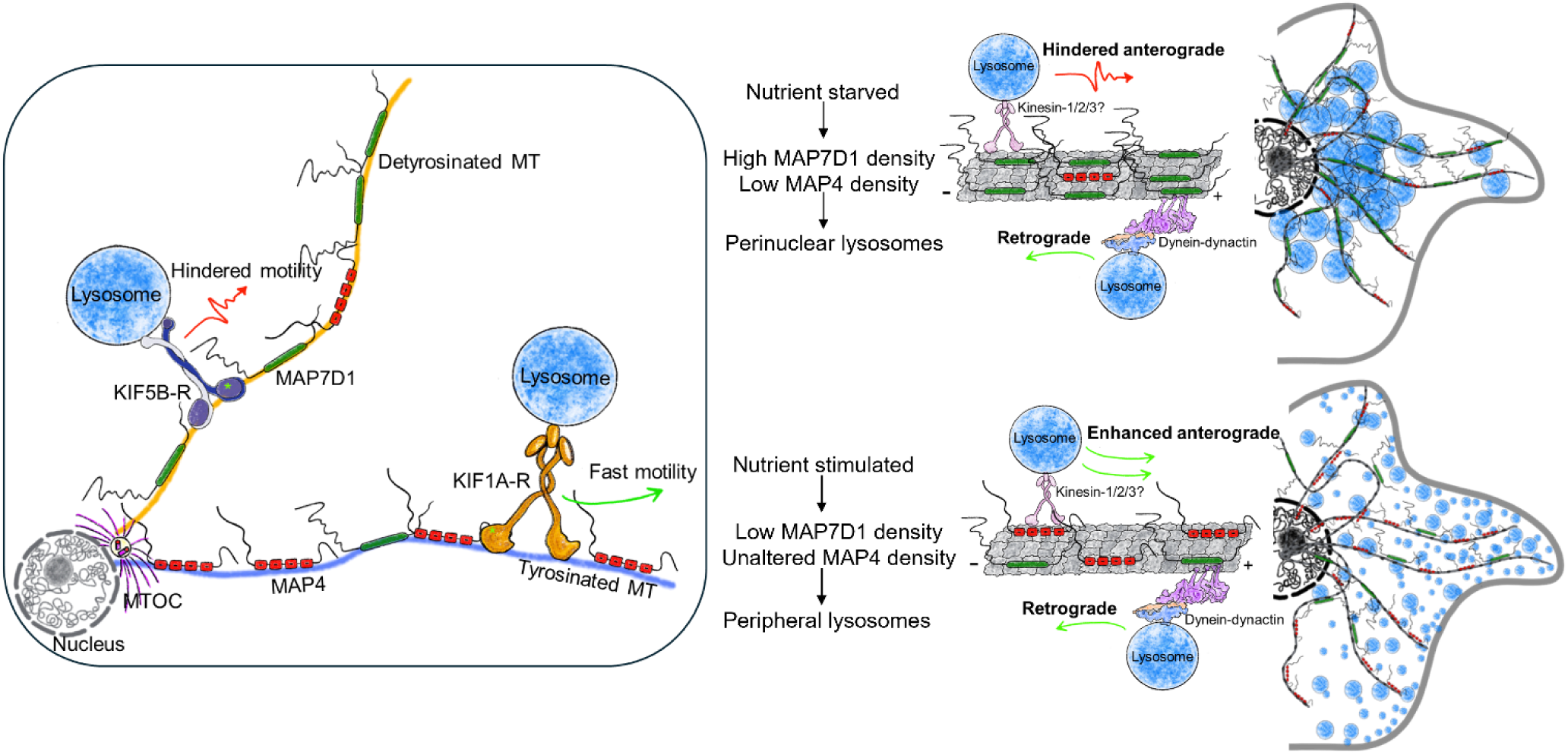
Proposed model The schematic presents a model integrating findings from this study and prior reports^6,27,29,30^. Under homeostatic conditions, the microtubule network is differentially decorated with MAPs: MAP4 is enriched on tyrosinated microtubules, while MAP7D1 is enriched on detyrosinated microtubules. These MAP–PTM combinations establish specialized tracks that spatiotemporally regulate lysosome transport. On MAP7D1-decorated detyrosinated microtubules, kinesin-1 is preferentially recruited but its motility is dampened by the high density of MAP7D1 nanoclusters, promoting lysosomal immobilization in the perinuclear region to support autophagy-related functions. In contrast, MAP4-enriched tyrosinated microtubules favor fast kinesin-3 or kinesin-2–driven anterograde transport, redistributing lysosomes toward the cell periphery to enhance mTORC1 signaling. Importantly, anterograde transport is dynamically balanced by dynein-driven retrograde movement, which counteracts kinesin activity and maintains equilibrium between perinuclear and peripheral lysosome pools during homeostasis. Upon nutrient modulation, MAP densities are remodelled along microtubule tracks. Nutrient deprivation increases MAP7D1 density and reduces MAP4 abundance, thereby immobilizing kinesin-1 and biasing transport toward dynein-driven retrograde motility, which retains lysosomes in the perinuclear region. Conversely, nutrient stimulation decreases MAP7D1 density, while maintaining optimal MAP4 density, thereby enabling faster motility of kinesin and promoting anterograde transport and peripheral positioning of lysosomes. This adaptive tuning of MAP organization in response to nutrient availability provides a mechanism for spatial reorganization of lysosomes to meet cellular metabolic demands.

A previous study noted that MAP4 is depleted from microtubule bundles and cellular extensions enriched in detyrosinated tubulin^40^, but whether this heterogeneity reflected MAP4’s selective binding remained unclear due to limitations in imaging resolution. Using super-resolution microscopy, we now demonstrate that MAP4 preferentially associates with tyrosinated microtubules and is excluded from detyrosinated ones, independent of acetylation status. In contrast, MAP7, previously linked to acetylated microtubules^37,39^, now preferentially decorates acetylated-detyrosinated microtubules and sparsely associates with acetylated-tyrosinated filaments, further refining the microtubule code for MAP7 association. Together, these findings reveal a spatial paradigm: tyrosination and detyrosination segregate distinct MAPs, while acetylation distributes MAP gradients across the network.

MAP selectivity could be coupled with microtubule lattice conformation. Recent studies show that MAPs can modulate lattice spacing, with MAP7 and CAMSAPs promoting expansion^37,38^. This expansion facilitates access for α-TAT1 and VASH1–SVBP, consistent with MAP7’s enrichment on acetylated-detyrosinated microtubules. Furthermore, short-term Taxol-induced expansion redistributed MAP7 onto tyrosinated microtubules. We anticipate that with prolonged Taxol exposure, these MAP7-decorated tyrosinated microtubules would progressively acquire acetylation or detyrosination, reflecting enhanced substrate accessibility of the PTM-enzyme on an expanded lattice.

The facilitation of detyrosination on expanded lattices raises the question of whether MAP4 restricts detyrosination by compacting the lattice. However, unlike other tau-family MAPs such as Tau or MAP2, MAP4 neither compacts the lattice nor dissociates upon expansion^44^. Consistent with this, MAP4 localization on tyrosinated microtubules remained unchanged following Taxol-induced expansion. These observations suggest that MAP4 exclusion from detyrosinated filaments is not lattice-driven but likely mediated through direct inhibition of VASH. A recent study demonstrated that MAP4 restricts VASH activity in a concentration-dependent manner^36^. Supporting this, MAP4 overexpression reduced detyrosination, while a projection-domain-deleted MAP4 mutant neither restricted detyrosination nor avoided binding to detyrosinated microtubules. These findings suggest a critical role for the projection domain in MAP4’s selective association with tyrosinated microtubules. Though classically linked to microtubule bundling^46^, projection domains are increasingly recognized for their roles in modulating motor-driven transport^47^. We propose that the projection domain stabilizes MAP4 interaction with tyrosinated microtubules or mediates steric interference with VASH or other factors associated with detyrosinated microtubules. Given their intrinsically disordered nature, elucidating how the projection domains mechanistically confer microtubule binding specificity to MAP4 remains challenging^32^.

Functionally, MAP gradients across PTM-defined microtubules create regulatory zones that govern motor activity and organelle dynamics. MAP4 has been shown to inhibit kinesin-1 (KIF5B) while promoting kinesin-3 (KIF1A) motility in a concentration-dependent manner^32,44^. Consistent with this, our data show that MAP4-decorated tyrosinated microtubules localize kinesin-3 (KIF1A-R) and exclude kinesin-1 (KIF5B-R). Similarly, Tau—a neuronal MAP closely related to MAP4—exhibits differential inhibition of motor proteins, requiring ∼10-fold higher concentrations to inhibit dynein than kinesin, suggesting directional control of transport^33^. Our data demonstrates that high MAP4 levels impair anterograde lysosome motility over retrograde transport, while optimal MAP4 concentrations on tyrosinated microtubule support fast long-range transport toward cell membrane enabling peripheral lysosome positioning to promote mTORC1 activation. Consistent with our findings, recent studies have shown that MAP4 can spatially activate PI3K-Akt signaling on endosomes by interacting with PI3KC2α, which acts upstream of and synergizes with mTORC1 signaling^48,49^.

Conversely, MAP7 family members are known to facilitate kinesin-1 recruitment and activation while inhibiting kinesin-3^29–31^. Consistently, we show that MAP7-enriched detyrosinated microtubules localize kinesin-1 (KIF5B-R) and exclude kinesin-3 (KIF1A-R). Here we show that near saturating concentration of MAP7D1 on microtubule renders lysosome immobile while lower concentrations were permissive of faster motility. Our observations are consistent with in vitro studies that shows MAP7 modulates kinesin-1 motility bi-phasically, promoting motility at optimal levels but inhibiting it at near-saturating concentrations^29^. Additionally, our prior work demonstrated that kinesin-1-driven lysosomal transport is inhibited on detyrosinated tracks, facilitating autophagosome–lysosome fusion^27^. Together these evidence suggests that differential enrichment of MAP7D1 on detyrosinated microtubule promotes perinuclear accumulation of lysosomes to enable autophagosome-lysosome fusion in a kinesin-1 dependent manner.

This model also explains the nutrient-dependent remodeling of MAP densities enabling lysosome positioning and function. During starvation, MAP7D1 density increases, thereby inhibiting kinesin-1 motility and thus immobilizing lysosomes in the perinuclear region.

Further, sub-optimal density of MAP4 during starvation might not facilitate kinesin-3 mediated anterograde lysosome movement and relieve the inhibition on dynein, thereby tipping the balance towards retrograde transport. In contrast, during nutrient stimulation MAP7D1 density decreases, which activates kinesin-1 and MAP4 density remains optimal to promote kinesin-3. Thus kinesin-1/2/3 alone or in combination mediate the anterograde transport promoting the peripheral lysosome positioning. Thus, MAP densities are adaptively tuned in response to nutrient cues to orchestrate lysosomal organization. Together, these findings outline a functionally adaptive molecular toolbox: MAP4–kinesin–3–tyrosinated microtubules support peripheral lysosome distribution to promote mTORC1 signalling, while MAP7D1–kinesin–1– detyrosinated microtubules promote perinuclear accumulation of lysosomes to enable autophagy (Figure 9).

Importantly, *in vitro* reconstitution experiments show that MAP4 does not inherently distinguish between tyrosinated and detyrosinated microtubules, and kinesin-1 motility is not directly influenced by microtubule acetylation or detyrosination (REF). These contrasts highlight the role of the cellular environment in shaping MAP–PTM–motor interactions. Within cells, the local concentrations and spatial organization of MAPs, PTM enzymes, and motors constitute a dynamic molecular toolbox tuned to physiological needs. Supporting this, recent work has shown that osmotic stress triggers cytoskeletal remodelling to enhance MAP7 association and microtubule acetylation, thereby biasing exocytosis-endocytosis dynamics to maintain cell size and homeostasis^37^. By analogy, we propose that the MAP4–kinesin–3– tyrosination and MAP7–kinesin-1–detyrosination axes operate as context-sensitive circuits, selectively engaged in response to nutrient cues to orchestrate lysosomal dynamics and signalling.

What remains unresolved are the cellular mechanisms by which MAP densities on microtubules are modulated in response to nutrient signaling. Possible pathways include (i) phosphorylation of MAPs by nutrient-sensitive kinases to regulate association with microtubules^50^, (ii) alterations in microtubule lattice state by other MAPs that drive redistribution^37,38^, (iii) modulation of PTM levels to favor or disfavor MAP–PTM interactions, or changes in microtubule dynamic instability. Future studies will be required to determine how these mechanisms converge to remodel MAP organizations in response to cellular signals.

## Materials and Methods

### Cell Culture and Transient Transfection

African green monkey (Cercopithecus aethiops) kidney epithelial cells (BS-C-1; a kind gift from Prof. Melike Lakadamyali, University of Pennsylvania, USA) and Human cervical carcinoma (HeLa) cells were maintained in a complete growth medium, i.e. MEM (GIBCO #10370021) and DMEM (GIBCO #11995065), respectively. The medium was supplemented with nonessential amino acids, 10% (v/v) fetal bovine serum (FBS) (Hyclone FBS, #SH30071.03), 2 mM L-glutamine (GIBCO #25030081), 1 mM sodium pyruvate (GIBCO #15140122), and penicillin-streptomycin (GIBCO #15140122) or AntiAnti (GIBCO #15240062). Cell culture condition was maintained at 37°C with 5% carbon dioxide. The cell lines were routinely checked for mycoplasma contamination using the Mycoplasma PCR Detection Kit (Abcam, ab289834).

For experiments, the cells were seeded on either #1.5 coverslip bottom 35mm confocal dishes (Ibidi, Gräfelfing, Germany) or eight-well chambers (Ibidi, Gräfelfing, Germany) for imaging. Transient transfection of desired plasmids was performed in cells using X-tremeGENE™ HP DNA Transfection Reagent (Roche) for 24 – 48 hours. A complete list of plasmids used in this study is provided in #Table 1.

**Table #1.** List of Plasmids used in the study.

| **Table#1** |  |
| --- | --- |
| **Plasmid** | **Source** |
| mEGFP-C1 | Addgene #54759 |
| mCherry2-C1 | Addgene #54563 |
| pGEMHE-mEGFP-Map4 | Addgene #105571 |
| mRuby-MAP4-C-10 | Addgene #55873 |
| pEGFP-N1-TFEB | Addgene #38119 |
| pEGFP-Map7-full-length | Addgene #46076 |
| eGFP-KIF5B-Rigor | Addgene #172204 |
| eGFP-KIF1A-Rigor | Addgene #172207 |
| GFP-TTL | Kind gift from Prof Minhaj Sirajuddin and Prof. Carsten Janke |
| GFP-VASH2 | Kind gift from Prof Minhaj Sirajuddin and Prof. Carsten Janke |
| KIF5B-Rigor-mCherry | This paper |
| MAP4-FL-mCherry | This paper |
| MAP4-FL-EGFP | This paper |
| MAP4-MTBD | This paper |
| MAP4-MTBD-ΔN | This paper |
| MAP4-MTBD-ΔC | This paper |

**Table #2.** List of Primary Antibodies used in the study.

| **Table #2** |  |  |  |
| --- | --- | --- | --- |
| **Antibodies** | **Host Species** | **Identifier** | **Supplier** |
| Anti-MAP4 | Rabbit | ab224543 | Abcam |
| Anti-MAP4 | Mouse | sc-390286 | Santa Cruz Biotechnology, Inc |
| Anti-Tyrosinated tubulin | Rat | MAB1864-I | EMD Millipore |
| Anti-Detyrosinated tubulin | Rabbit | ab48389 | Abcam |
| Anti-acetylated tubulin | Mouse | T6793 | Sigma Aldrich |
| Anti-LAMP2 | Mouse | sc-18822 | Santa Cruz Biotechnology, Inc |
| Anti-MAP7 antibody | Mouse | H00009053-B01P | Abnova |
| Anti MAP7D1 antibody | Rabbit | HPA028075 | Sigma-Aldrich/Atlas |

### Treatments

#### Lysotracker deep red treatment

For labelling lysosomes using LysoTracker deep red dye, a 1 µM stock solution of LysoTracker deep red dye was diluted to a final working concentration of 125 nM for labelling lysosomes. Subsequently, 250 µL of this working solution was added to the cells for 15 min at 37°C in the dark.

#### Taxol treatment

For the microtubule lattice expansion assay, cells were incubated with 100 µM Taxol (Sigma Aldrich #T7191) or Dimethyl sulfoxide (DMSO, Sigma Aldrich #D2650) in complete media for 10 minutes at 37°C. Following treatment, the cells were fixed with ice-cold methanol and rinsed with PBS before further processing for immunostaining.

#### Nutrient starvation and stimulation treatment

BS-C-1 cells were starved by incubating them for 3 hours in Krebs-Ringer’s Buffer solution (KRBH, pH 7.4) supplemented with 4.5 mM glucose, 0.1% bovine serum albumin, and 1 mM sodium pyruvate. For nutrient stimulation, control cells were incubated in 2x amino acid medium—complete MEM media, supplemented with 2× MEM amino acids (R, C, H, I, L, K, M, F, T, W, Y, V; MEM amino acids ×50 liquid, GIBCO 11130051) for 20 minutes^51,52^.

#### siRNA Treatment

ON-TARGETplus SMARTpool siRNA against human MAP4 (L-011724-01-0005) was purchased from Dharmacon. For scramble siRNA, select Negative Control No. 2 siRNA was purchased from (Thermo Fisher Scientific #4390846). The siRNA was transfected using X-tremeGENE™ siRNA Transfection Reagent (Roche) at final siRNA concentrations of 50 to 100 nM.

#### Generation of Plasmids

mEGFP-Full-length MAP4 was obtained from pGEMHE-mEGFP-Map4 (Addgene Plasmid #105571 from Melina Schuh) by digestion with EcoRI and XmaI and cloned into mEGFP-C1 vector backbone (Addgene Plasmid #54759 from Michael Davidson). Microtubule Binding Domain (MTBD) of MAP4 mRuby-MAP4-C-10 (Addgene Plasmid #55873 from Michael Davidson).

MAP4-MTBD domain was subcloned into the mEGFP-C1 backbone using the following primers:

5′-TCAGAATTCATCTGGTGGCTCCGCAGGATCC-3′ (forward) and

5′-ACAGGTACCCTATCCCACTTTGGCTTGGGC-3′ (reverse);

MAP4-N-MTBD domain was subcloned from mEGFP-Full-length MAP4 using the following primers:

5′-TAATGATGACATTCTGGTCTCGTCCGC-3′ (forward) and

5′-TCCCACTTTGGCTTGGGCCTT-3′ (reverse);

MAP4-MTBD-C domain was subcloned from mEGFP-Full-length MAP4 using the following primers:

5′-TCCCGGCAAGAAGAAGCAAAG-3′ (forward) and

5′-CATTCTGTCGACTGCAGAATTCGA-3′ (reverse);

The ATPase “rigor” mutants of Kinesin-1 (Addgene Plasmid #172204 eGFP-KIF5B-Rigor (G234A) and Kinesin-3 (Addgene Plasmid #172207 eGFP-KIF1A-Rigor (G251A) Juan Bonifacino. The KIF5B-Rigor was cloned into mCherry2-C1 (Addgene Plasmid #54563 from Michael Davidson) using 5′-CGGGGTACCAATTCGAATTCTATGGCGGAC-3′ (forward) and

5′-AATGGATCCTTACGACTGCTTGCCTCCA-3′ (reverse).

#### Immunofluorescence

For immunofluorescence, the cells were first rinsed with 1x PBS pH7.4 (GIBCO #10010023). Fixation was done either with absolute Methanol (Sigma Aldrich) for 7 minutes at −20°C or 4% Paraformaldehyde (Electron Microscopy Sciences, #15710) in PBS for 10 minutes at room temperature (RT). Post fixation, cells were washed with PBS and blocked at room temperature for 2 hours in blocking buffer (3% w/v BSA in 0.2% Saponin (w/v; Sigma Aldrich #47036). Cells were then incubated with primary antibodies diluted at the appropriate ratio in blocking buffer overnight at 4°C. A complete list of antibodies used in this work is provided in Table #2. Subsequently, cells were incubated with dye-conjugated secondary antibodies for 1 hour at room temperature. Washing steps between antibody incubation were performed with 1x PBS pH 7.4 three times each.

### TIRF and STORM Imaging

TIRF and STORM imaging were performed using a custom-built microscope setup based on a Nikon Ti2E microscope body with a 100× oil immersion TIRF objective (NA 1.49; Nikon). The system has a quad-band filter set (TRF89902-ET-405/488/561/647; Chroma Technology) and an EM-CCD camera (iXon Ultra 897, Andor Technology). For TIRF imaging, appropriate laser lines were used to excite fluorophores and fluorescence emission was collected through the same objective. Images were captured at an exposure time of 100 ms per frame. For live-cell imaging, 1000 frames were acquired, while 10 frames per field of view were acquired for fixed-cell imaging. For 2D STORM imaging, the same TIRF setup was used with laser excitation at 647 nm (Coherent or MPB Communications) to excite Alexa Fluor 647 dye (Invitrogen #A20006), and a 405-nm solid-state laser (Obis; Coherent) to reactivate the dye via the Alexa Fluor 405 activator dye (Invitrogen # A30000). STORM image acquisition was performed at 20 ms exposure time per frame, with 60,000 frames captured per image. STORM images were analyzed and rendered as previously described using custom-written software, Insight3, a kind gift from Prof. Bo Huang, University of California, San Francisco, CA^53^.

### Correlative imaging and single particle tracking

For correlative imaging of lysosomes and endogenous MAP4 STORM, the live imaging of LAMP1-YFP vesicles was performed on the TIRF microscope using a 488 nm laser, as described above. After imaging, cells were fixed on the stage and processed for immunostaining endogenous MAP4, as mentioned previously. The super-resolution STORM images of MAP4 in the same cell were captured as explained above.

To correct for potential sample drift during the fixation and immunostaining procedures, we aligned the live lysosome channel with the STORM image of MAP4 as described previously^27,54^. We coated the imaging dishes with fiduciary markers (0.1 μm TetraSpeck beads; Invitrogen #T7279) before seeding the cells. The beads were visible during the live and STORM imaging, and the mean position of multiple beads was estimated by localizing the beads repeatedly during single particle tracking and STORM analysis. The shifts calculated in x and y were then applied to the raw localization files of the STORM, and alignment was validated by comparing bead positions before and after correction.

For single particle tracking of lysosomes, a custom-developed, semi-automated particle tracking software Track Multiple and a custom MATLAB script as described previously^27,54^. The x and y coordinates were obtained by fitting a 2D Gaussian function to the point spread function of the lysosomes. Further analysis to identify direct and confined phases was conducted using a custom MATLAB-based script, based on a moving window approach, using four-point segments along the data. For each segment, a ratio of the straight-line displacement between the segment’s starting and ending points to the sum of stepwise displacements within the segment was computed. Since segments overlapped, individual trajectory points contributed to multiple segments, and the ratio for each point was averaged across all its associated segments. This ratio served as a measure of linearity, with values near 1 indicating active transport. A threshold linearity ratio of 0.7 was used to classify active and passive phases. To refine the classification further, we used an angle-based criterion: segments where consecutive displacement vectors formed angles smaller than 90° were classified as passive. Additionally, mean square displacement (MSD) analysis was used to validate the classification, where active transport exhibited an α value greater than 1.5. Only trajectories with at least five data points were considered for further analysis. Metrics such as the speed of direct phases and run phase (run length per unit track length) were calculated from the processed data.

### Confocal Imaging

Confocal images were acquired using a Zeiss laser scanning confocal microscope (LSM 780 system) equipped with a 63x oil immersion objective, a 488 nm Argon laser, a 561 nm DPSS laser and a 633 nm HeNe laser.

### SIM Imaging

3D Super-Resolution Structured Illumination Microscopy (3D SR-SIM) was performed using a completely motorized Zeiss Elyra 7 (Lattice SIM technology - Carl Zeiss Ltd). Images were acquired using a plan Apo 63×/1.40 oil objective and sCMOS camera (PCO Edge) with immersol 518F (23°C) immersion oil. Lattice SIM acquisition mode captured 15 phase images at 1,280 × 1,280-pixel resolution, and SIM processing was performed using ZEN Black (ZEISS) software to generate the final super-resolved image. All representative SIM images in the figures were adjusted for brightness and contrast using ZEN Lite software from Zeiss.

### Single-Particle Tracking (SPT) in TrackMate

To analyse the bulk motility parameters like net displacement we performed single particle tracking analysis of lysotracker-positive vesicles, cells were treated with LysoTracker™ Deep Red for endogenous and MAP4-FL overexpression. Following treatment, cells were imaged using TIRF microscopy. Imaging was performed at a 100 ms time interval for 100 sec. Raw data for lysotracker vesicle motility were processed by background subtraction using a rolling ball radius of 50.0, followed by bleach correction through histogram matching. The processed data were then analyzed using the TrackMate plugin^55^ in Fiji software with the following parameters:

- Vesicle diameter: 0.6 µm
- Detector: LoG (Laplacian of Gaussian)
- Initial thresholding: None
- Tracker: Simple LAP tracker
- Linking max distance: 1.3 µm
- Gap-closing max distance: 2 µm
- Gap-closing max frame gap: 20
- Filter on track: Number of spots in the track
- Mobile immobile population threshold on net displacement

### Estimation of the relative abundance of MAPs from SIM images

To quantify the mean intensity of endogenous MAP4 and MAP7D1 on different microtubule subsets (tyrosinated microtubules, tyrosinated-acetylated microtubules, and detyrosinated-acetylated microtubules), z-stack SIM images were processed into maximum intensity projections using Fiji. A fixed-size region of interest (ROI) (25 × 55 pixels) was selected using the rotated rectangle tool to define MAP-enriched regions on each microtubule subset. The mean intensity of MAPs within each ROI was measured using the “Measure” tool. To generate a pie chart representing the percentage of MAP on each microtubule subset, the mean intensity values from an equal number of ROIs for each subset were normalized to the maximum intensity within the dataset. Each value was then scaled to 100%, and the sum of all respective values within each category was used to represent the relative abundance of MAPs on each microtubule subset.

### Colocalization analysis

For colocalization analysis, Mander’s coefficient (M1 or M2) was determined using the JACoP plugin^56^ of Fiji software^57^.

### Estimation of Localization density

The localization density of MAP4 and MAP7D1 was quantified as the number of localizations per unit area of the defined region of interest.

### Cluster analysis and estimation of cluster parameters

Segmentation of localizations in STORM images to perform clustering was performed using the Hierarchical density-based clustering algorithm (HDBSCAN). A threshold of a minimum of 4 localizations and a minimum cluster size of 30 localizations was used as a parameter to segment the clusters. The elbow plot method was used to determine the epsilon in an unbiased manner. For cluster analysis, the ROI (region of interest) containing a single microtubule was cropped to ensure MAP4 clusters on adjacent microtubules do not contribute to further quantification. To estimate the size of MAP4 clusters identified through HDBSCAN analysis, the centre of mass of each cluster was calculated by taking the geometric mean of all localizations within the cluster. The outer boundary of each cluster was then approximated using a convex hull, which connects the farthest localizations within the cluster. Subsequently, the distance from the centre of mass to the convex hull boundary points was measured for each cluster, and the average of these distances was approximated as the cluster radius. This method provides a robust estimation of cluster size by incorporating both the centre of mass and edge boundaries defined by the spatial distribution of localizations. The centroid of adjacent clusters was connected, and the distance between the boundary of clusters along the line connecting the centroid was calculated as the inter-cluster distance. The number of clusters per micron of microtubule length was calculated as ratio of total number of MAP4 clusters per unit length.

### Percentage of MAP Localization density calculation

To quantify the relative abundance of MAP4 and MAP7D1 across microtubule segments, we analyzed manually segmented equal sized regions of interest (ROIs) across multiple cells. Within each ROI, the number of localizations corresponding to MAP4 and MAP7D1 were estimated separately. The localization percentage for each MAP was then calculated as the ratio

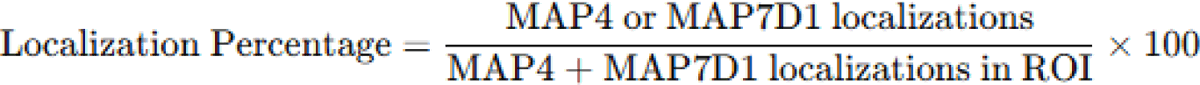

of MAP-specific localizations to the total localizations of both MAPs within that ROI:

The percentage localization densities of the two MAPs were plotted as stacked column graphs, and several ROIs from multiple images were sorted in increasing order of MAP4 localization percentage to visualize the trend and correlation between association of the two MAPs in each ROI.

### MAP4–MAP7D1 Overlap Analysis

To assess the degree of spatial overlap between MAP4 and MAP7D1 nanoclusters, we segmented the MAP clusters using HDBSCAN (as described earlier) for each ROI. Convex hulls were generated around each identified to define the spatial boundary of the cluster. All pairwise combinations of overlapping clusters between MAP4 and MAP7D1 were then evaluated for the extent of overlap. When an intersection between convex hull boundaries was detected, a binary mask was generated representing the shared region. Further we quantified the % contribution of localizations of each MAP in the overlapping region in terms of number of localizations within the overlapping region.

### Estimation of lysosome distribution

Confocal images were analyzed to assess the distribution of LAMP2-positive vesicles using the “Radial Profile” plugin in Fiji. To quantify lysosomal distribution, a circular region was drawn around the entire cell with the nucleus as the centre. The normalized integrated intensity of LAMP2 was then measured within this region. Data are presented as the mean lysosomal distribution ± SEM, obtained from multiple cells across three independent experiments.

### Estimation of the size and number of lysosomes

To estimate the area and number of vesicles (LAMP2-positive) from confocal images, each z-stack was converted to an 8-bit maximum intensity projection using the program. The area and number of vesicles were measured using the “Analyze Particle” Tool.

### Statistical Analysis

The data was exported to Microsoft Excel (2013) and Origin 2022b (Academic) for further analysis.

## Data and code availability statement

The raw data presented is accessible from the corresponding author (N.M.) upon request. The custom analysis codes written in Python are available as well.

## Acknowledgements

We thank Prof. Melike Lakadamyali (University of Pennsylvania, Philadelphia, PA) for supporting the research with reagents, software, critical reading of the manuscript and intellectual inputs. We thank Prof Bo Huang (University of California, San Francisco) for the STORM analysis software. We thank Prof. Appu Kumar Singh (IIT Kanpur) for insights on structural analysis. NM thanks Prof. Arun Kumar Shukla (IIT Kanpur) for suggestions and mentorship. D.M.K expresses gratitude towards the Prime Minister’s Research Fellowship (PMRF) program, India, for providing financial assistance during research. N.M. acknowledges the Ramalingaswami Re-entry Fellowship (BT/RLF/Re-entry/39/2018) of the Department of Biotechnology (DBT), India and Anusandhan National Research Foundation (ANRF)-Science and Engineering Research Board (SERB), India (CRG/2022/008391) for the funding and the Indian Institute of Technology (IIT), Kanpur, India, for infrastructure. The authors thank all lab members for their support. N.M acknowledges Mrs. Shefali Tripathi for her kind assistance during the initial stages of setting up the research.

## Author contribution

D.M.K. designed and carried out the experiments and performed the data analysis. J.K. performed the colocalization analysis and quantification of microtubule-detyrosination levels. S.B. and S.C.C. wrote the code for cluster analysis. D.M.K and N.M. designed the research and wrote the manuscript. N.M. conceptualised, supervised, and acquired funding for the research. All authors provided feedback on the manuscript.

