## Supplementary Material for "MAP4-MAP7D1 partitioning on tyrosinated-detyrosinated microtubules coordinates lysosome positioning in nutrient signalling"

### Supporting Information:

#### Figure S1:

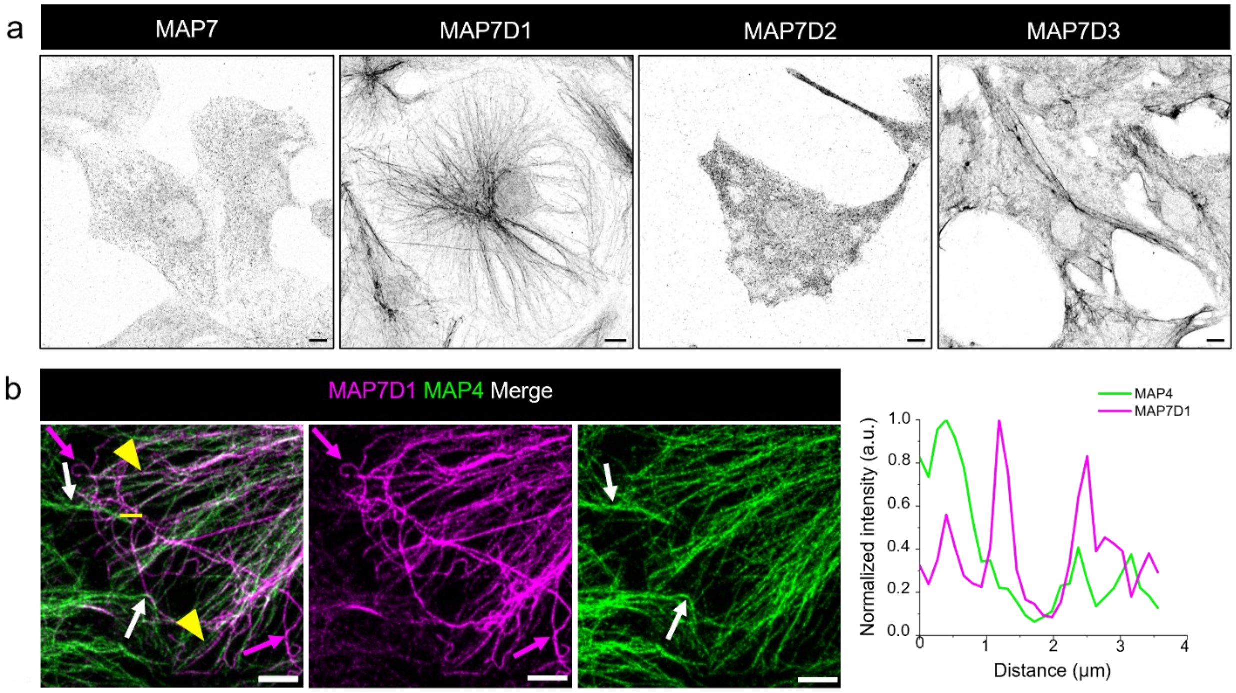


**Figure S1. MAP4 and MAP7D1 localizes to distinct microtubules in BS-C-1 cells.**
(a). Representative confocal images for endogenous levels of four MAP7 isoforms in BS-C-1 cells highlight that MAP7-D1 is the predominant microtubule-bound isoform present in the BS-C-1 cell line. (b). Representative two-color confocal images of endogenous MAP7D1 (magenta) and MAP4 (green) in BS-C-1 cells. To ensure that the observed minimal overlap in figure 1A (MAP7D1 first followed by MAP4) was not due to antibody competition, sequential immunostaining was performed (MAP4 first, followed by MAP7D1. Magenta arrows indicate MAP7D1-dominated microtubules, white arrows indicating MAP4-dominated microtubules, and yellow arrowheads marking microtubules decorated by both MAPs. A corresponding line intensity plot along the yellow line in the inset reveals that the two MAPs are predominantly associated with distinct microtubule tracks. Scale bars: 10 μm for (a); 5 μm for (b).

### Figure S2:

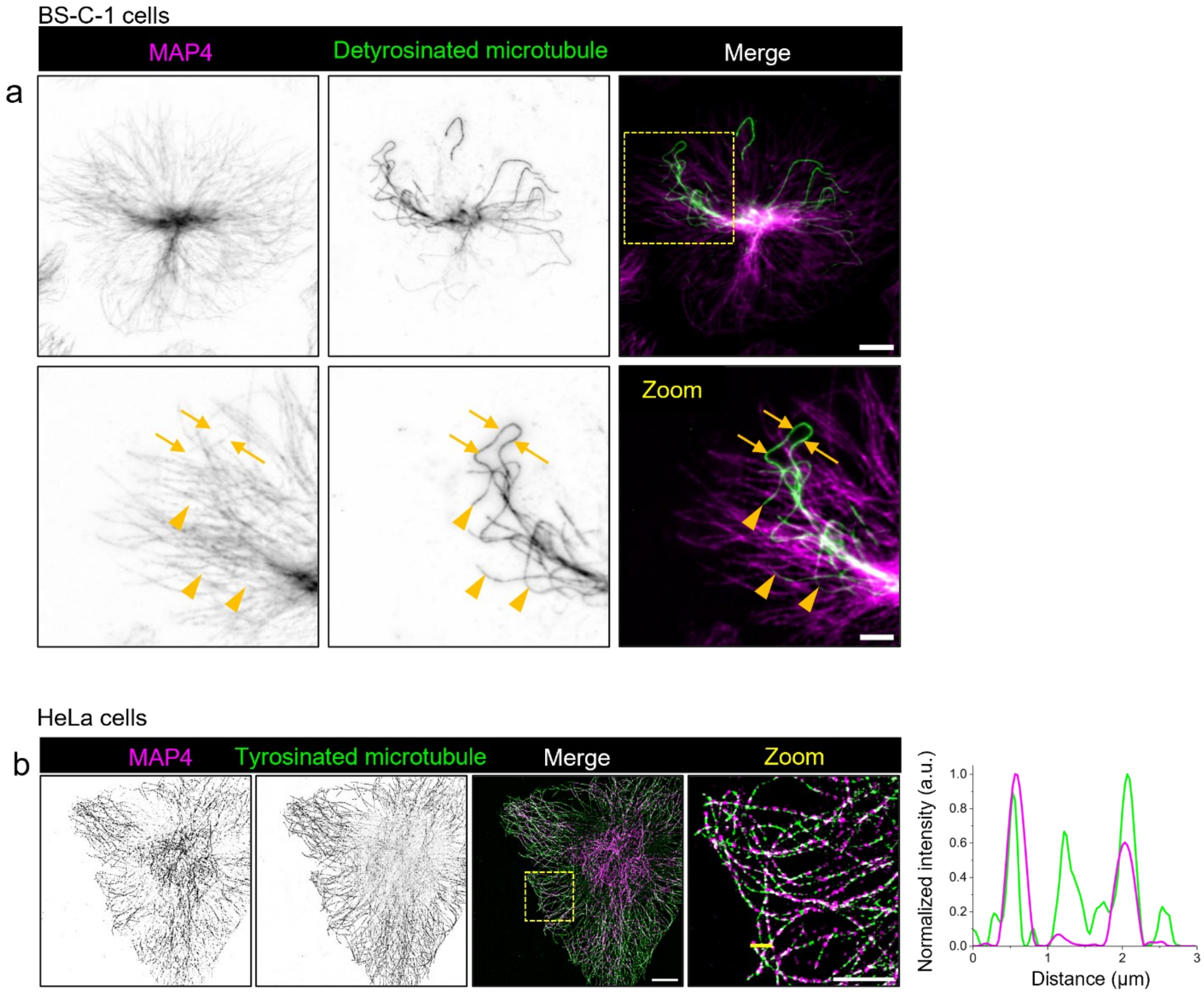


**Figure S2. MAP4 selectively localizes to tyrosinated microtubules.**

(a) TIRF images (grayscale inverted) of endogenous MAP4 (magenta) with detyrosinated microtubules (green) in BS-C-1 cells. Zoomed inset (yellow box) shows that most detyrosinated microtubules either completely lack MAP4 (arrowheads) or minimally decorated by MAP4 (arrows). (b) SIM images of endogenous MAP4 and tyrosinated microtubules in HeLa cells, along with corresponding line intensity plots, demonstrate that MAP4's preferential binding to tyrosinated microtubules is conserved across epithelial cell lines. Scale bars: 10 μm for (a, b); 5 μm for zoomed images.

### Figure S3:

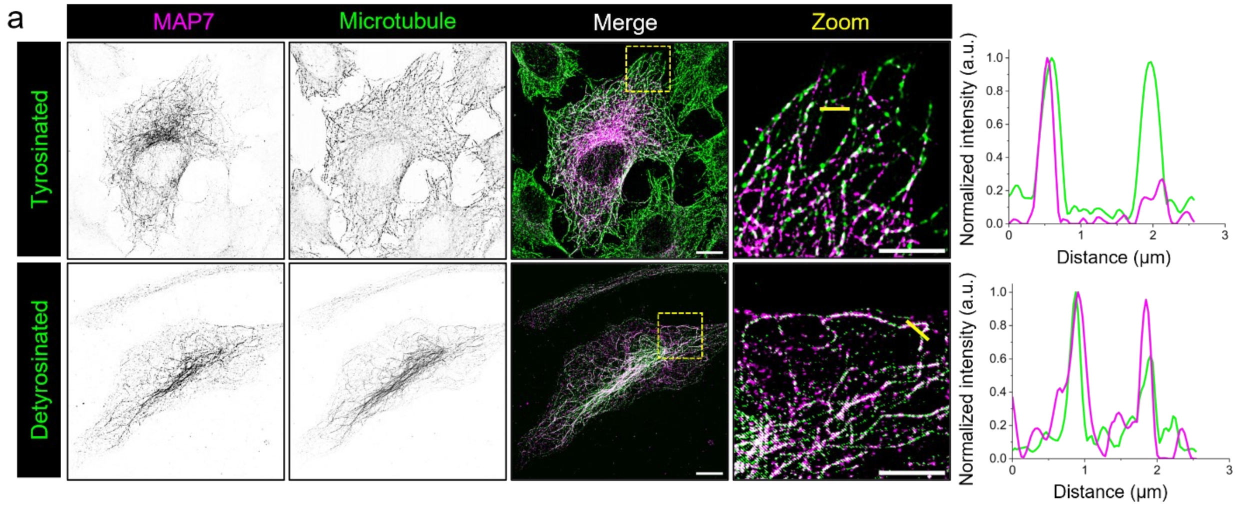


**Figure S3. MAP7 differentially localizes to tyrosinated and detyrosinated microtubules.**

(a) Super resolution SIM images of endogenous MAP7 (magenta) with detyrosinated microtubules (green) in HeLa cells. Corresponding line intensity plots along yellow line in zoomed image show MAP7 localizes to both linear tyrosinated and curved detyrosinated microtubules, albeit with a higher preference for the detyrosinated microtubule subset. Scale bars: 10 μm for image and 5 μm for zoomed images.

### Figure S4:

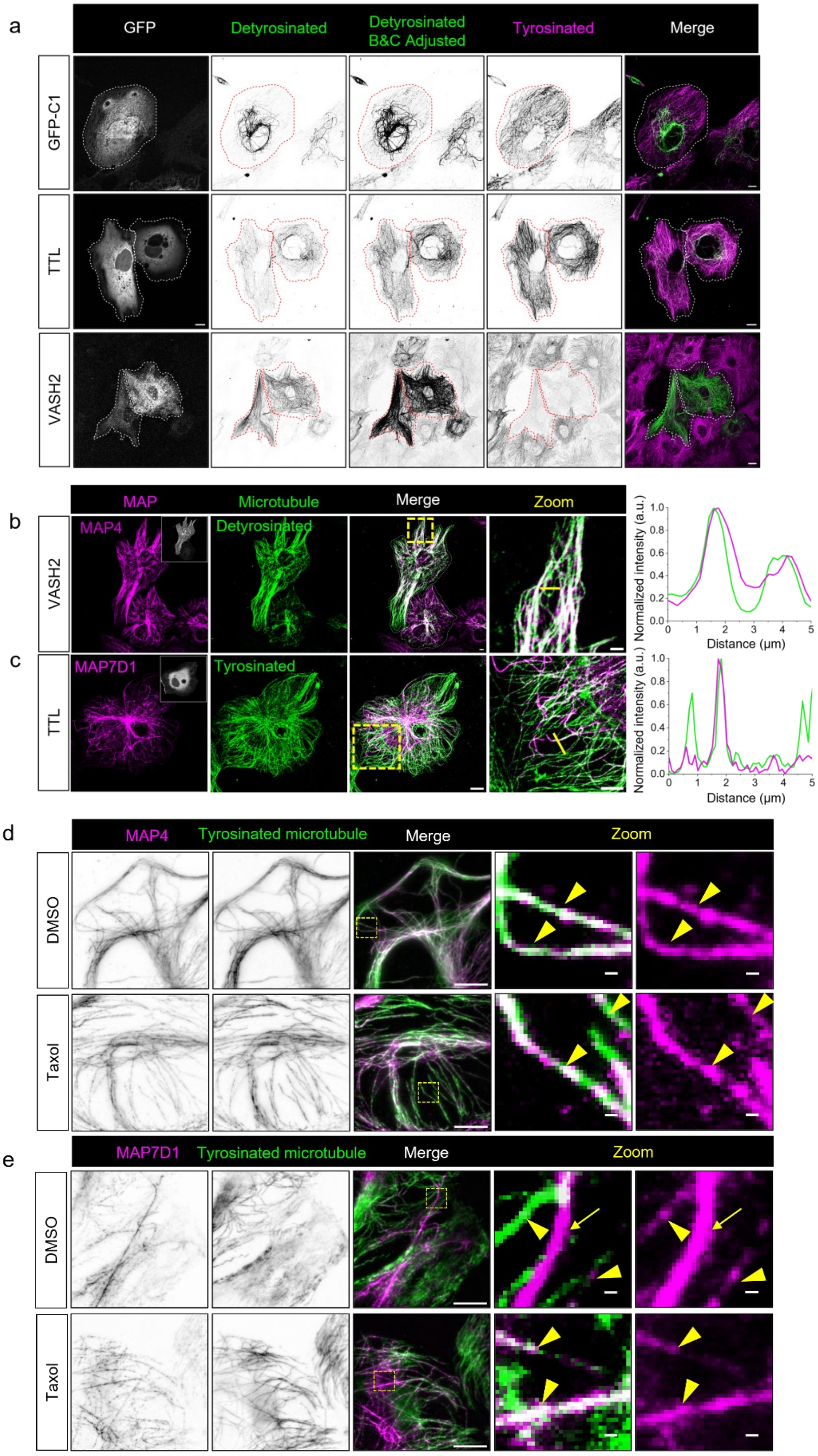

a
GFP
Detyrosinated
Detyrosinated B&C Adjusted
Tyrosinated
Merge
GFP-C1
TTL
VASH2
b
MAP
Microtubule
Merge
Zoom
MAP4
Detyrosinated
VASH2
Normalized intensity (a.u.)
Distance (μm)
c
MAP7D1
Tyrosinated
TTL
Normalized intensity (a.u.)
Distance (μm)
d
MAP4
Tyrosinated microtubule
Merge
Zoom
DMSO
Taxol
e
MAP7D1
Tyrosinated microtubule
Merge
Zoom
DMSO
Taxol

**Figure S4: Modulation of microtubule-PTMs by enzyme overexpression and lattice expansion by Taxol.**

(a) Confocal images of BS-C-1 cells transiently expressing GFP vector control, TTL-GFP, or VASH2-GFP (GFP signal in white), immunostained for tyrosinated (magenta) and detyrosinated (green) microtubules. TTL expression markedly reduced overall percentage of detyrosinated microtubules relative to tyrosinated ones, whereas VASH2 expression significantly increased them. (b) Representative confocal images and corresponding line intensity profiles of BS-C-1 cells transiently expressing GFP-VASH2 inset, white) show endogenous MAP4 (magenta) localizing extensively on detyrosinated microtubules (green). (c) Cells expressing GFP-TTL (inset, grey) reveal that endogenous MAP7D1 (magenta) remains enriched on a subset of microtubules that are largely tyrosinated (green). (d-e) Two-color TIRF images showing endogenous MAP4 (magenta) or MAP7D1 (magenta) with tyrosinated microtubules (green) in DMSO or Taxol-treated cells. ROIs were drawn on tyrosinated microtubules overlapping with respective MAP signals. Yellow arrowheads marks MAP on tyrosinated microtubule and arrows indicate not on tyrosinated microtubule. These ROIs were used to quantify MAP4 or MAP7D1 localization density from their corresponding STORM images (see figure 3g, h) in DMSO versus taxol-treated conditions. Scale bars: 10 µm (full image a–e), 2µm (insets b, c), 500nm (insets d, e).

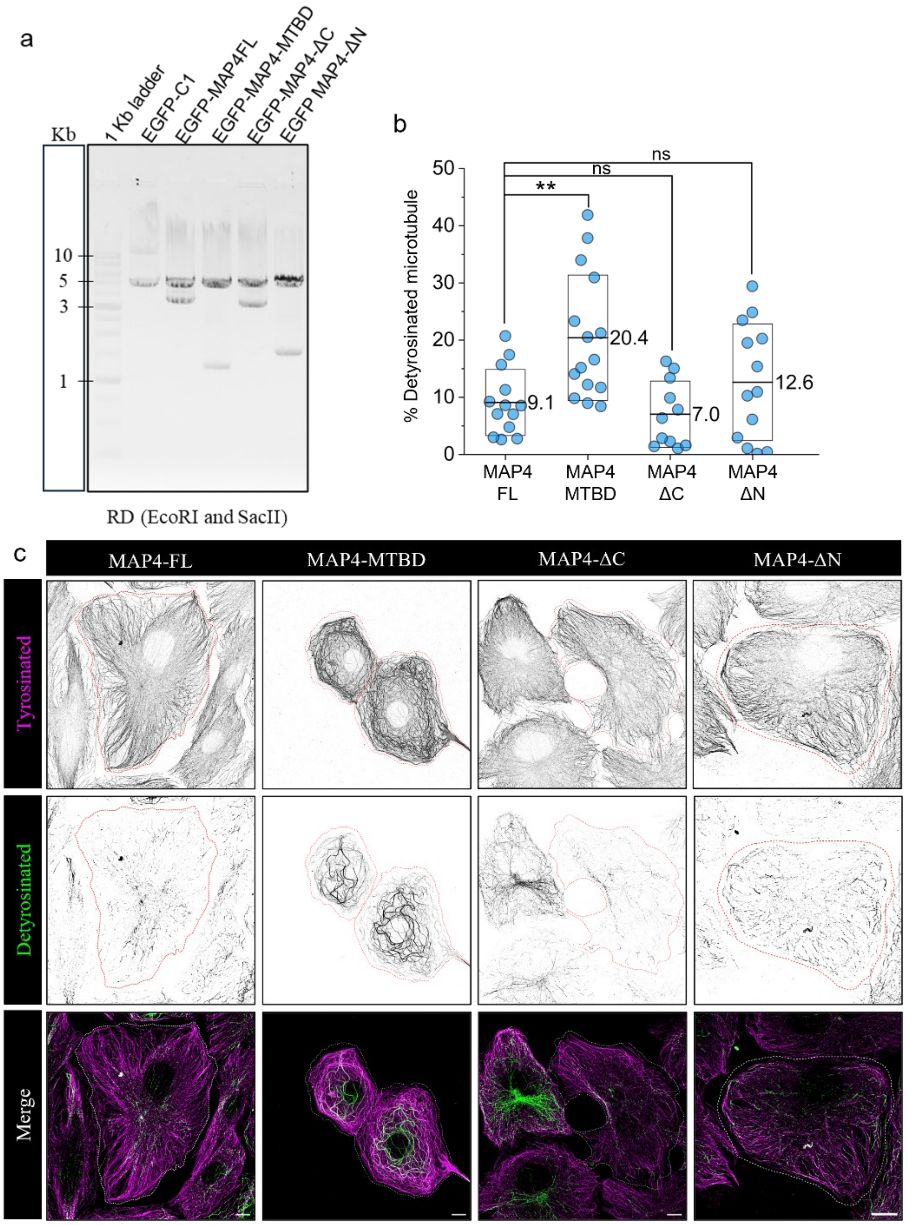


**Figure S5: MAP4 projection domain regulates microtubule detyrosination.**

(a) Verification of MAP4 deletion constructs (generated in this study) using agarose gel electrophoresis following restriction digestion with EcoRI and SacII. Insert size differences reflect specific truncations. (b) Box plot quantifying % Detyrosinated microtubule in cells expressing indicated constructs. (c) Representative confocal images (grayscale inverted) of BS-C-1 cells transiently expressing EGFP-MAP4 variants immunostained for tyrosinated (magenta) and detyrosinated (green) microtubules. Transfected cells in images are highlighted by marking the boundaries. Data represent mean (line) ± SD (box). Number of cells (n) analysed for MAP4-FL (13), MAP4-MTBD (15), MAP4-ΔN (13) and MAP4-ΔC (14). Statistical significance was determined using the Mann–Whitney U test (**$p<0.01$, not significant [ns]). Scale bars: 10 μm

### Figure S6:

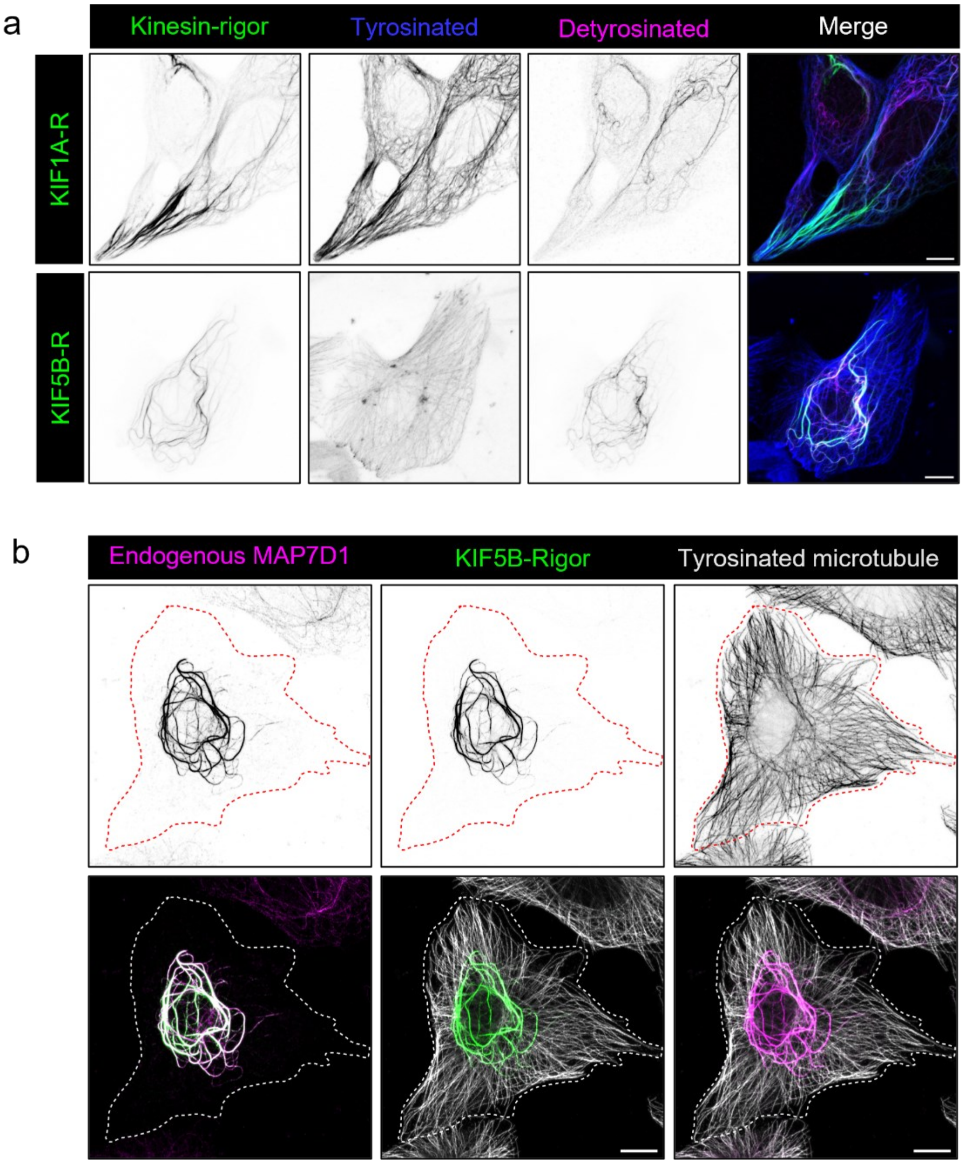


**Figure S6. KIF1A and KIF5B rigor mutants are associated with distinct microtubule subsets.**

(a) Confocal images of BS-C-1 cells transiently expressing GFP-tagged (a) KIF1A-R (green) or (b) KIF5B-R (green) immunostained for tyrosinated (blue) and detyrosinated (magenta) microtubules, highlighting that KIF5B-R and KIF1A-R predominantly decorate detyrosinated and tyrosinated microtubules, respectively. (b) BS-C-1 cells expressing KIF5B-R (green), immunostained for endogenous MAP7D1 (magenta) and tyrosinated microtubules (white), showing that MAP7D1 is more concentrated on KIF5B-R decorated detyrosinated microtubules, and is almost depleted from tyrosinated microtubules in cells expressing KIF5B-R. Scale bars: 10 µm.

### Figure S7:

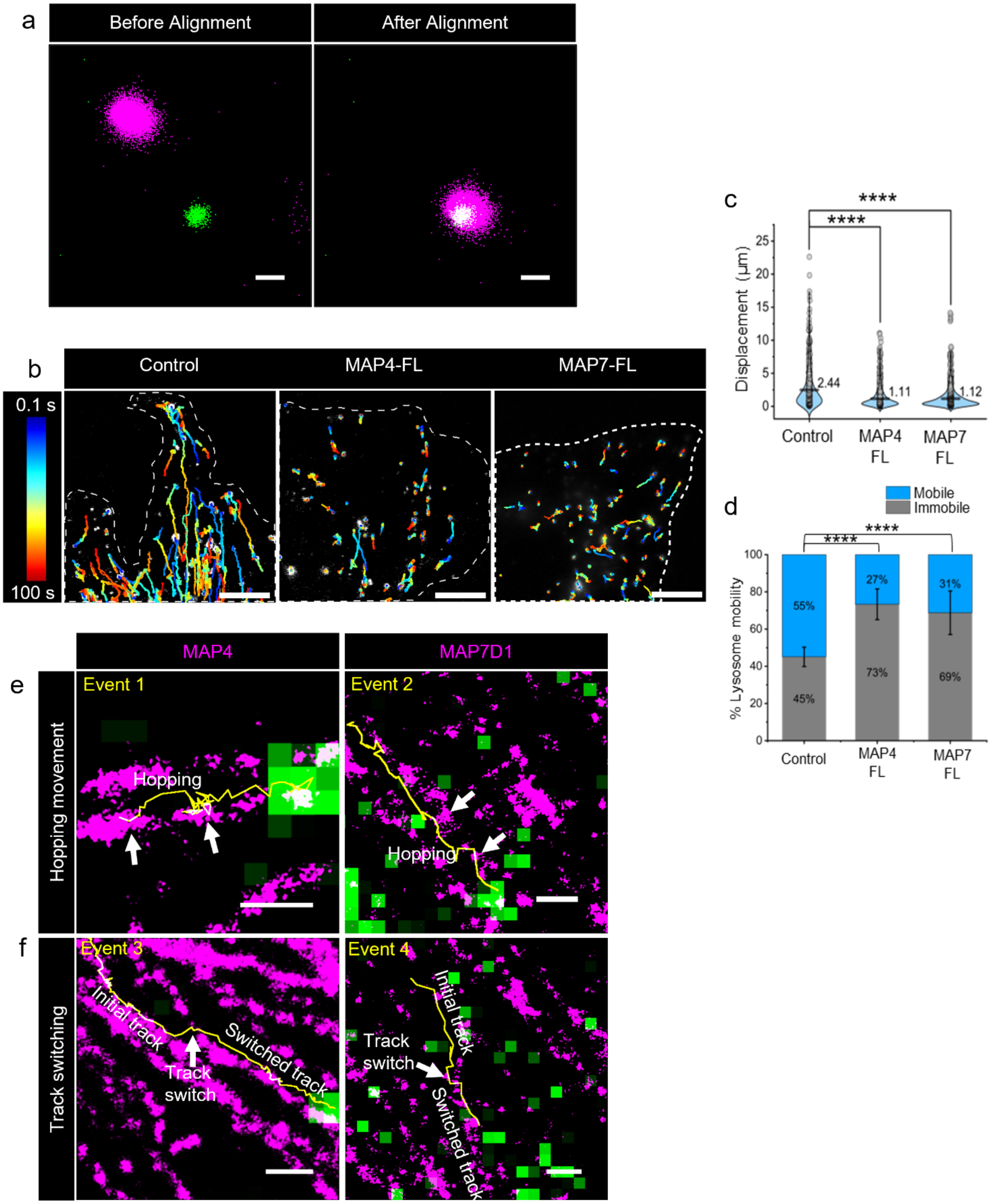

a
Before Alignment
After Alignment
b
Control
MAP4-FL
MAP7-FL
0.1 s
100 s
c
****
****
Displacement (μm)
25
20
15
10
5
0
2.44
1.11
1.12
Control
MAP4 FL
MAP7 FL
d
Mobile
Immobile
****
****
% Lysosome mobility
100
80
60
40
20
0
55%
45%
27%
73%
31%
69%
Control
MAP4 FL
MAP7 FL
MAP4
MAP7D1
e
Hopping movement
Event 1
Hopping
Event 2
Hopping
f
Track switching
Event 3
Initial track
Track switch
Switched track
Event 4
Initial track
Track switch
Switched track

**Figure S7: MAPs density on microtubule regulates lysosomal motility**

(a) Localizations corresponding to the same Tetraspeck beads imaged during live TIRF (green) and STORM (magenta) image acquisition used for image registration and alignment. Localizations from the same bead are shown before and after alignment. (b) Representative trajectories (time (t) colour-coded as shown in the inset) of lysotracker-positive vesicles from TrackMate analysis of live-cell TIRF imaging (captured at 10 fps for 1 min) in BS-C-1 cells with control (no. of cells = 9), MAP4-FL overexpression (no. of cells = 14) and MAP7-FL overexpression (no. of cells = 12). (c) Violin plot for net displacement of lysosomal trajectories in control (no. of lysosomal trajectories = 720), MAP4-FL overexpression (no. of lysosomal trajectories =1033), and MAP7-FL overexpression (no. of lysosomal trajectories =1599). The black line represents the mean. (d) Bar plot quantifying the percentage of mobile and immobile lysosomes in control, MAP4-FL and MAP7-FL overexpression (mean ± s.e.m.). Representative overlays of lysosomal trajectories (yellow) with MAP4 and MAP7D1 STORM images (magenta) illustrating distinct lysosomal trafficking behaviours. (e) Example of a trajectory hopping across MAP4 (event 1) or MAP7D1 (event 2) nanoclusters (white arrows). (f) Example of a track-switching event (white arrow) upon encountering MAP4 (event 3) or MAP7D1 (event 4) nanoclusters. The initial track and switched tracks are labelled respectively. Scale bars: (a) 100 nm; (b) 10 μm; (e, f): 500 nm.

### Figure S8:

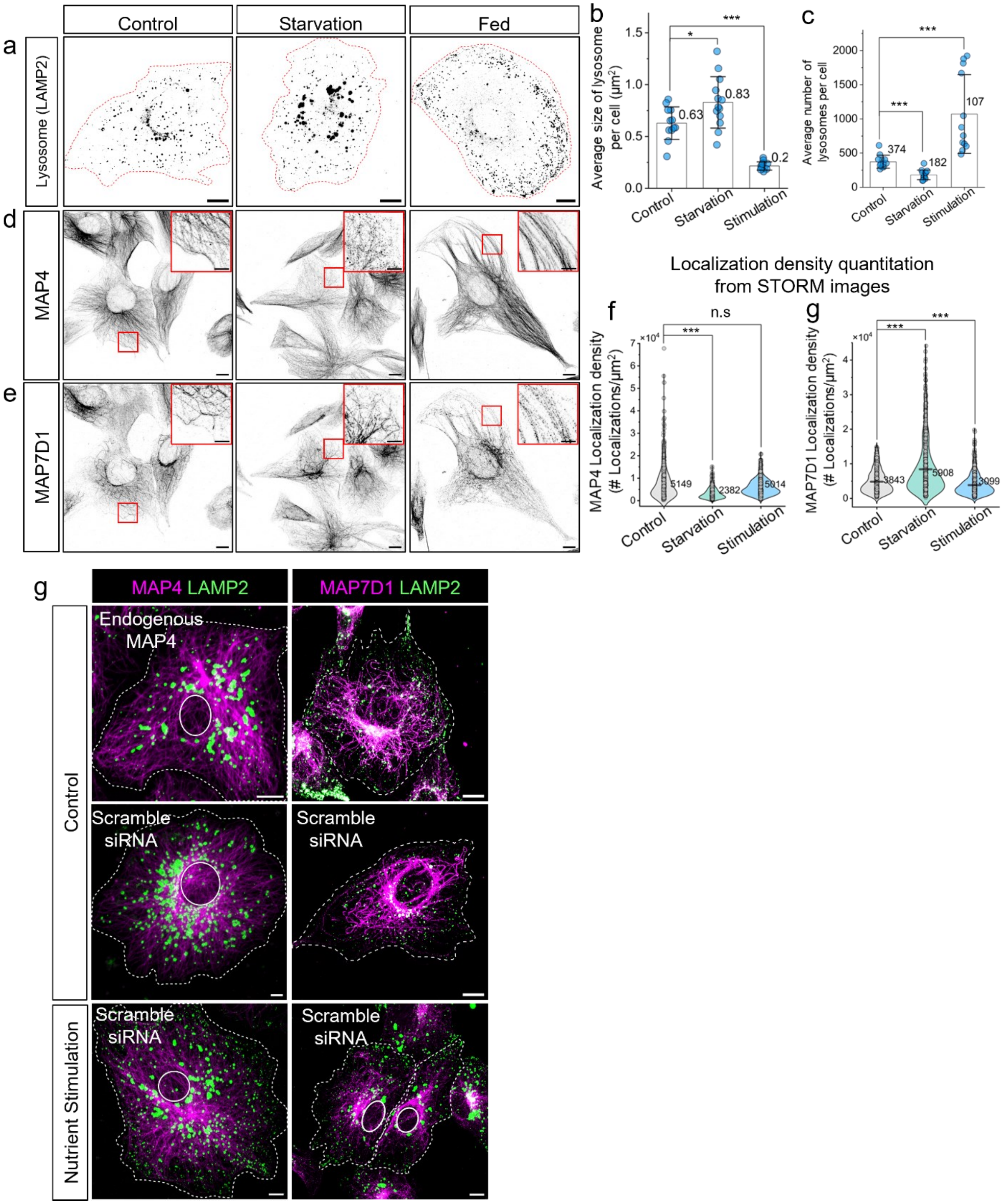

a
Control
Starvation
Fed
Lysosome (LAMP2)
b
Average size of lysosome per cell (μm²)
0.63
0.83
0.2
Control
Starvation
Stimulation
c
Average number of lysosomes per cell
374
182
107
Control
Starvation
Stimulation
d
MAP4
e
MAP7D1
Localization density quantitation from STORM images
f
n.s
MAP4 Localization density (# Localizations/μm²)
5149
2382
5014
Control
Starvation
Stimulation
g
MAP7D1 Localization density (# Localizations/μm²)
3843
5908
3099
Control
Starvation
Stimulation
g
MAP4 LAMP2
MAP7D1 LAMP2
Control
Endogenous MAP4
Scramble siRNA
Scramble siRNA
Nutrient Stimulation
Scramble siRNA
Scramble siRNA

**Figure S8. Cells tune MAP density on microtubules in response to nutrients.**

(a) Representative grayscale inverted confocal images of lysosomes (LAMP2) along with bar graph quantifying (b) average lysosome size per cell and (c) average lysosome number per cell under control (n=12), nutrient starvation (n=13) and 2X amino acid stimulation (n=12) conditions, highlighting modulation in position, size and number of lysosomes. (d-e) Representative inverted grayscale confocal images of endogenous MAP4 and MAP7D1 in the same cells under different nutrient conditions highlight striking differences in the density of MAPs on microtubules. (f) Violin plots for localization density of MAP4 and MAP7D1 (replotted from Figure 8d, f to highlight median values) under different nutrient conditions quantified from 3 different cells STORM images. (g) Representative dual colour confocal image of endogenous MAP4 or MAP7D1 (magenta) with lysosomes (green) in control or nutrient-stimulated cells. Scale bar: 10 μm, insets in d, e are 5 μm.

**Supporting video legends:**

#### Movie S1

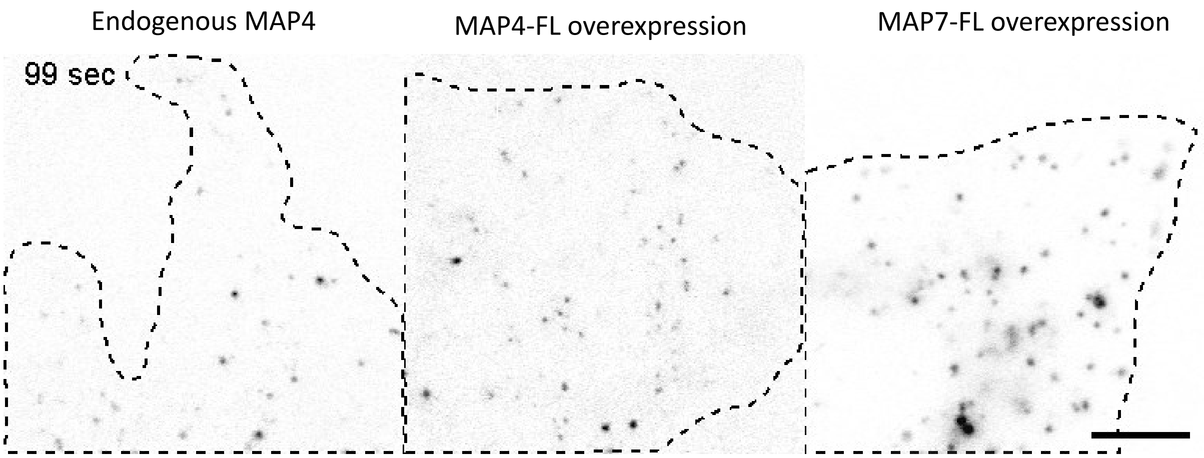


**Movie S1.** Live-Cell imaging of lysotracker positive vesicles under Control and transiently expressing MAP4-FL, and MAP7-FL. Related to Figure S7b. Cells were recorded at 10 fps for 100 s. MAP4-FL and MAP7-FL overexpression leads to a marked increase in the immobile population of lysotracker-positive vesicles as compared to control cells. Speed 10x, Scale bar: 10 µm.

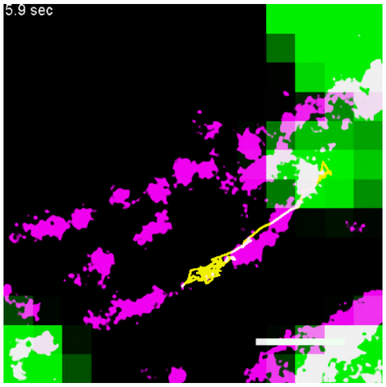


#### Movie S2

**Movie S2.** Correlative live-cell imaging of lysosome (green) moving in anterograde direction (yellow trajectory) overlaid with STORM image of endogenous MAP4 (magenta). Related to Figure 7a. Cells were recorded at 10 fps. Speed 1x, Scale bar: 500 nm.

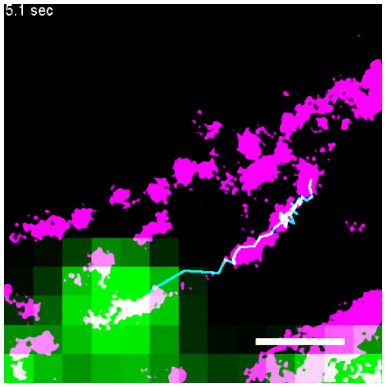


#### Movie S3

**Movie S3.** Correlative live-cell imaging of lysosome (green) moving in retrograde direction (cyan trajectory) overlaid with STORM image of endogenous MAP4 (magenta). Related to Figure 7a. Cells were recorded at 10 fps. Speed 1x, Scale bar: 500 nm.

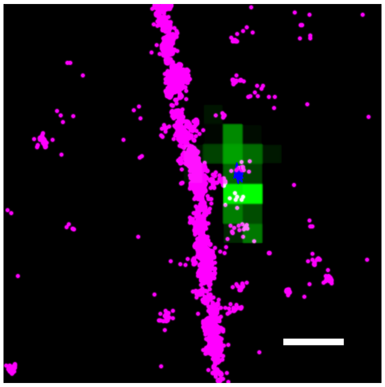

#### Movie S4

**Movie S4.** Correlative live-cell imaging of lysosome (green) immobile (blue trajectory) overlaid with STORM image of endogenous MAP7D1 (magenta) at very high density. Related to Figure 7e. Cells were recorded at 10 fps. Speed 1x, Scale bar: 500 nm.

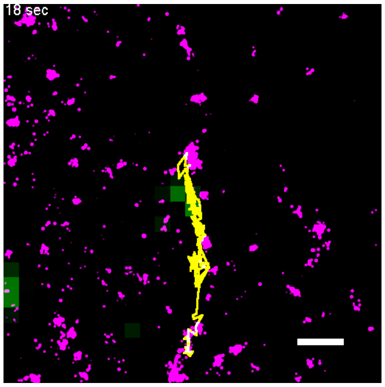


#### Movie S5

**Movie S5.** Correlative live-cell imaging of lysosome (green) moving in anterograde direction (yellow trajectory) overlaid with STORM image of endogenous MAP7D1 (magenta). Related to Figure 7e. Cells were recorded at 10 fps. Speed 1x, Scale bar: 500 nm.

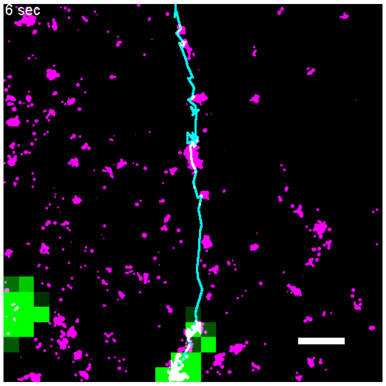


#### Movie S6

**Movie S6.** Correlative live-cell imaging of lysosome (green) moving in retrograde direction (cyan trajectory) overlaid with STORM image of endogenous MAP7D1 (magenta). Related to Figure 7e. Cells were recorded at 10 fps. Speed 1x, Scale bar: 500 nm.

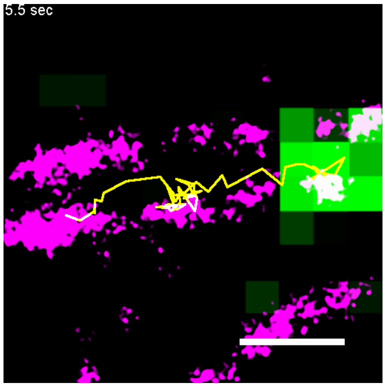


#### Movie S7

**Movie S7.** Correlative live-cell imaging of lysosome (green) hopping across MAP4 nanocluster (magenta), followed by a pause and pass behaviour. Related to Figure S7e. Cells were recorded at 10 fps. Speed 1x, Scale bar: 500 nm.

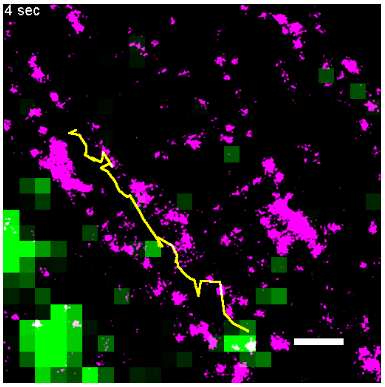


#### Movie S8

**Movie S8.** Correlative live-cell imaging of lysosome (green) hopping across MAP7D1 nanocluster (magenta), followed by a pass behaviour. Related to Figure S7e. Cells were recorded at 10 fps. Speed 1x, Scale bar: 500 nm.

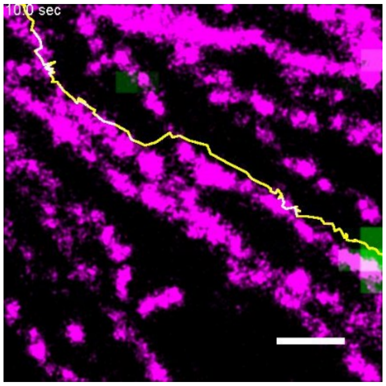


#### Movie S9

**Movie S9.** Correlative live-cell imaging of lysosome (green) switches track onto the adjacent microtubule at the junction of MAP4 nanocluster (magenta), followed by a movement on the same track. Related to Figure S7f. Cells were recorded at 10 fps. Speed 1x, Scale bar: 500 nm.

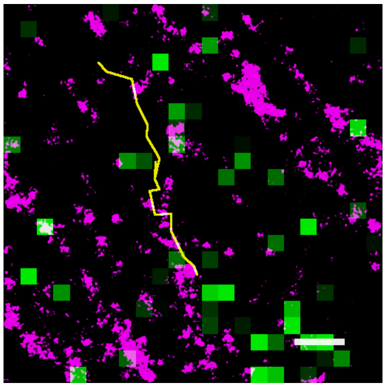

#### Movie S10

**Movie S10.** Correlative live-cell imaging of lysosome (green) switches track onto the adjacent microtubule at the junction of MAP7 nanocluster (magenta), followed by a movement on the same track. Related to Figure S7f. Cells were recorded at 10 fps. Speed 1x, Scale bar: 500 nm.
